# Beneficial effects of cerebellar tDCS on motor learning are associated with altered putamen-cerebellar connectivity: a simultaneous tDCS-fMRI study

**DOI:** 10.1101/2020.08.11.246850

**Authors:** Matthias Liebrand, Anke Karabanov, Daria Antonenko, Agnes Flöel, Hartwig R. Siebner, Joseph Classen, Ulrike M. Krämer, Elinor Tzvi

**Affiliations:** Department of Neurology, University of Lübeck, Lübeck, Germany; Danish Research Center for Magnetic Resonance, Centre for Functional and Diagnostic Imaging and Research, Copenhagen University Hospital Hvidovre, Hvidovre, Denmark; Department of Neurology, University Hospital Greifswald, Greifswald, Germany; German Centre for Neurodegenerative Diseases (DZNE) Standort Greifswald, Greifswald, Germany; Department of Psychology, University of Lübeck, Lübeck, Germany; Department of Neurology, University of Leipzig, Leipzig, Germany

## Abstract

Non-invasive transcranial stimulation of cerebellum and primary motor cortex (M1) has been shown to enhance motor learning. However, the mechanisms by which stimulation improves learning remain largely unknown. Here, we sought to shed light on the neural correlates of transcranial direct current stimulation (tDCS) during motor learning by simultaneously recording functional magnetic resonance imaging (fMRI). We found that right cerebellar tDCS, but not left M1 tDCS, led to enhanced sequence learning in the serial reaction time task. Performance was also improved following cerebellar tDCS compared to sham in a sequence production task, reflecting superior training effects persisting into the post-training period. These behavioral effects were accompanied by increased learning-specific activity in right M1, left cerebellum lobule VI, left inferior frontal gyrus and right inferior parietal lobule during cerebellar tDCS compared to sham. Despite the lack of group-level changes comparing left M1 tDCS to sham, activity increase in right M1, supplementary motor area, and bilateral superior frontal cortex, under M1 tDCS, was associated with better sequence performance. This suggests that lack of group effects in M1 tDCS relate to inter-individual variability in learning-related activation patterns. We further investigated how tDCS modulates effective connectivity in the cortico-striato-cerebellar learning network. Using dynamic causal modelling, we found altered connectivity patterns during both M1 and cerebellar tDCS when compared to sham. Specifically, during cerebellar tDCS, negative modulation of a connection from putamen to cerebellum was decreased for sequence learning only, effectively leading to decreased inhibition of the cerebellum. These results show specific effects of cerebellar tDCS on functional activity and connectivity in the motor learning network and may facilitate the optimization of motor rehabilitation involving cerebellar non-invasive stimulation.

## 1. Introduction

Learning motor skills is basic for our daily routines, as for instance when learning to ride a bicycle, or to play the piano. Imaging studies show that multiple cortical and subcortical structures are involved in this process (Hardwick et al., 2013), probably as part of a network mediating the different phases of learning (Doyon et al., 2009). Learning of a new motor sequence typically follows three phases: an early stage when learning is rapid and improvement is taking place within minutes of practice, a late stage when performance becomes automatic and improvements are smaller and only observable after several hours of practice, and a consolidation stage in which further gains in performance are evident without any practice. A theoretical model of motor learning suggests a cortico-striato-cerebellar network operating differently at each of these stages (Doyon et al., 2009). In the early stage a wide network consisting of striatum, cerebellum, motor cortical regions, prefrontal cortex and parietal cortex is thought to be recruited to establish the new motor routine. During the late stage, the model suggests that a limited part of this network, namely a cortico-striatal loop, is recruited for learning of motor sequences.

In order to link specific brain structures to motor learning and to investigate tools that may enhance learning (Polanía et al., 2018), previous studies have used non-invasive brain stimulation techniques such as transcranial direct current stimulation (tDCS) (Buch et al., 2017). TDCS is thought to enable modulation of synaptic plasticity in the stimulated brain region and thus may have an impact on a wide range of behaviors including motor learning (Stagg and Nitsche, 2011). In a seminal work by (Nitsche and Paulus, 2000), anodal tDCS to M1 led to increased amplitudes of motor evoked potentials, a signature for cortical excitability, whereas cathodal tDCS led to reduced motor evoked potentials. Later studies showed that anodal tDCS of M1 during motor sequence learning, i.e. online, led to marked enhancement in learning, evident in progressive reduction of reaction times (RT) over the course of training the motor sequence (Kantak et al., 2012; Krause et al., 2016; Nitsche et al., 2003; Saucedo Marquez et al., 2013; Stagg et al., 2011; Waters-Metenier et al., 2014). Interestingly, learning enhancements were observed also when anodal tDCS to M1 was applied offline during consolidation of a motor sequence (Rumpf et al., 2017). This effect is however not specific to M1. Other studies showed that anodal stimulation of the cerebellum has a similar effect on sequence learning (Ferrucci et al., 2013; Shimizu et al., 2017), but see (Jongkees et al., 2019). In addition, cerebellar tDCS also leads to improvements in general motor skill learning, assessed with visuomotor tasks (Block and Celnik, 2013; Cantarero et al., 2015; Galea et al., 2011; Hardwick and Celnik, 2014; Herzfeld et al., 2014; Spampinato and Celnik, 2017), although some evidence suggests null or opposite effects (Jalali et al., 2017; Panouillères et al., 2015). Thus, it seems that tDCS provides a powerful tool to alter activity patterns in the motor learning network, most prominently in M1 and cerebellum, and therefore to study the underpinnings of motor learning.

What is then the exact role M1 and cerebellum play in the process of motor learning and as part of the motor learning network? It has been postulated that while the cerebellum is important for the early stage through error-based learning, M1 is important for retention of the motor memory during the late stage and in consolidation (Galea et al., 2011; Penhune and Doyon, 2005). To address how M1 and cerebellum interact with each other during the different learning phases, we previously used a functional magnetic resonance imaging (fMRI)-based network approach (dynamic causal modelling, DCM) while participants learned a motor sequence. These studies showed that in the early learning stage, M1 but also premotor cortex and putamen connections to cerebellum were negatively modulated by sequence learning, potentially leading to decreased cerebellar activity (Tzvi et al., 2015, 2014). In the current study, we combined fMRI with online anodal tDCS to further investigate how modulation of M1 and cerebellar activity using tDCS, affects connectivity patterns within the cortico-striato-cerebellar learning network.

Previous studies combining tDCS and fMRI or positron emission tomography (Antal et al., 2011; Zheng et al., 2011) demonstrated that anodal tDCS over M1 during motor performance leads to both localized activity increase and to activity modulation in other brain regions (Lang et al., 2005), suggesting an effect on network interactions. This idea is supported by resting state (RS) functional connectivity studies showing increased connectivity in the motor network following anodal tDCS of M1 (Amadi et al., 2014; Polania et al., 2012; Sehm et al., 2012). We therefore hypothesized that online anodal tDCS to M1 or cerebellum would lead to activity increase below the anodal electrode, as well as to increased activity in other regions of the motor learning network, specifically during sequence learning. In addition, we hypothesized that online anodal tDCS will lead to changes in connectivity patterns between M1 and cerebellum leading to enhancements in learning performance.

## 2. Materials and methods

### 2.1. Participants

Twenty-seven healthy subjects participated in the study. All participants were right-handed as assessed with a short form (Veale, 2014) of the Oldfield handedness inventory (Oldfield, 1971) and had normal or corrected to normal vision. None of the subjects reported history of neurological or psychiatric disorders, or regular practice of activities involving exercise of motor sequences (playing music instruments or playing on the computer). Informed written consent was given by the participants prior to study participation. The study was approved by the Ethics Committee of the University of Lübeck and was conducted in accordance with the Declaration of Helsinki. Two subjects (females, aged 21 and 26) were excluded from the study due to technical malfunctions, resulting in a sample of 25 subjects (15 females; mean age: 22.6 years; range: 18-31).

We determined the sample size based on our previous DCM-fMRI studies using this paradigm (Tzvi et al., 2017, 2015, 2014) with the same sample size. Importantly, based on previous tDCS studies using the SRTT, an effect size of 0.6 is expected (Ferrucci et al., 2013). At least 25 participants would be therefore required to find a difference in learning between tDCS and sham with α = 0.05 and power of 90%.

### 2.2. Experimental design

All participants completed three experimental sessions. These were kept at least 10 days apart to exclude possible aftereffects of stimulation on the following session. In each session, subjects received either anodal tDCS to right cerebellum (rCB tDCS), anodal tDCS to left M1 (lM1 tDCS) or sham tDCS with the stimulation location being counterbalanced between participants (Fig. 1C). During stimulation, participants performed a modified version of the serial reaction time task (SRTT, Nissen and Bullemer, 1987) in the MRI, thus tDCS was delivered online. Stimulation started when participants began the experiment. In each session, a different 8-element sequence was learnt to prevent cross-over effects and balanced across stimulation protocols. Participants were explicit about the presence of a sequence in the task. Each tDCS session lasted 20 min. The duration of tDCS matched approximately the length of the experimental task. Following stimulation and task-performance, participants were moved out of the scanner and after a short break performed in a completion task to assess explicit knowledge of the sequence they performed earlier (see description below).

**Figure 1.**
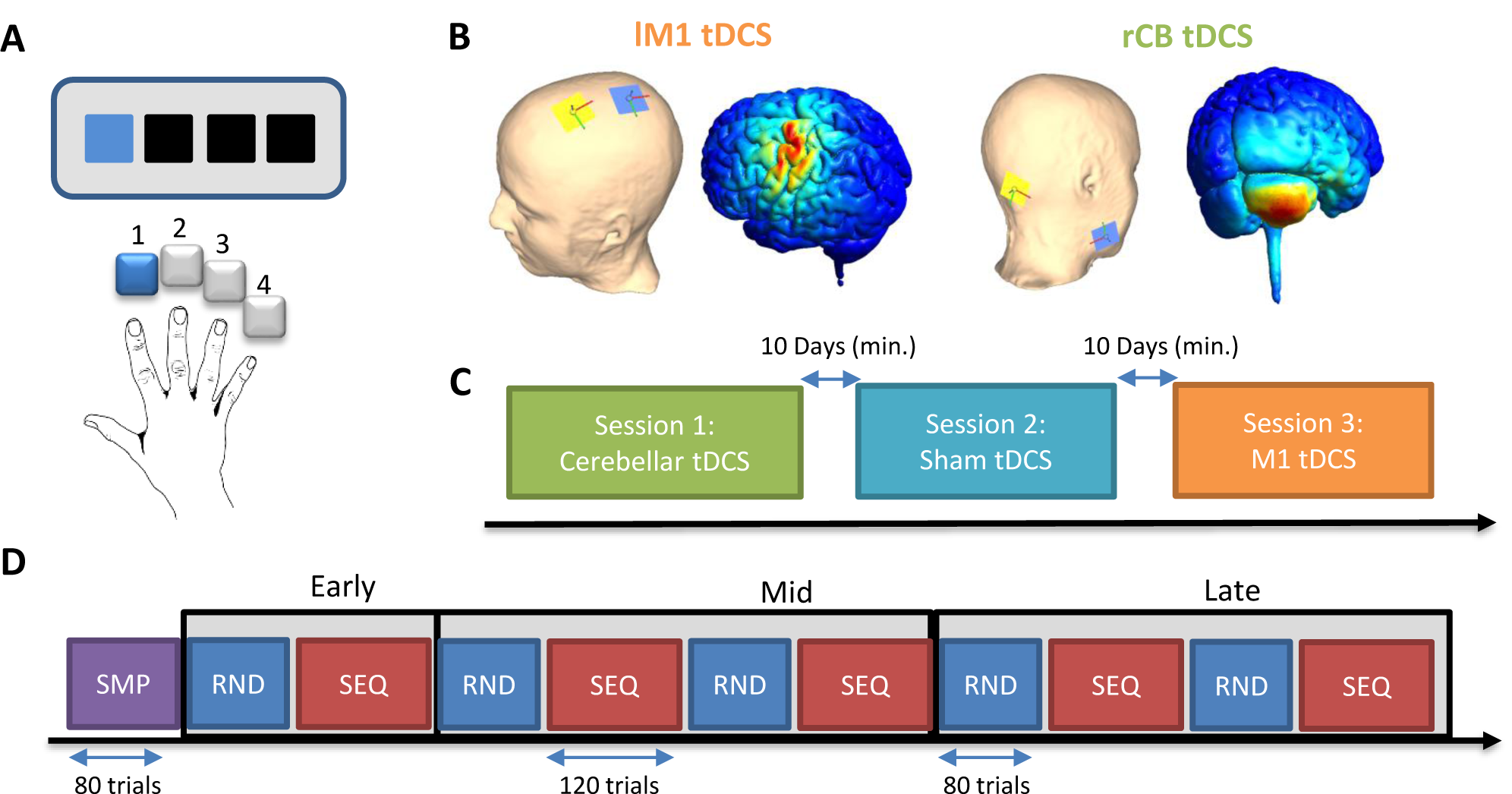
Experimental design. **A** Serial reaction time task. In each trial, 4 black squares were presented. At stimulus onset, one of the squares turned blue and subjects were instructed to press the button corresponding to the blue square with the respective finger. **B** Simulation of the current distribution for M1 montage (left) and cerebellar montage (right). On the dummy, a yellow electrode marks the location of the anodal electrode and blue of the cathodal electrode. **C** Experimental design. Subjects were invited to three sessions. In each session they received either cerebellar tDCS, M1 tDCS or sham. The order of sessions was randomized and counterbalanced for each subject. **D** In each session subjects performed the serial reaction time task in the MRI. Five random blocks (RND, blue) and 5 sequence blocks (SEQ, red) followed in an alternating order. A simple block (SMP, purple) appeared before or after the main task.

### 2.3. Serial reaction time task

In this task, participants were presented with four black squares on a grey colored background in a horizontal array, with each square (from left to right) associated with the following four fingers of the right hand: 1-index finger, 2-middle finger, 3-ring finger, 4-little finger (Fig. 1A). Participants were instructed to respond with the corresponding button as precisely and quickly as possible when one of the black squares turned blue. In case of a correct response, the blue square increased slightly in size. In case of a wrong response, the blue square turned red to mark the error and the next trial began. The response-stimulus interval was 500 ms. Visual stimuli were presented until the onset of a button press or the onset of the next trial. Trials were counted as correct when the appropriate key was pressed within 1000 ms after stimulus onset. In case no button was pressed within this time frame, a text appeared on the screen requesting the participants to be faster.

The task consisted of three different conditions: simple (SMP), random (RND), and sequence (SEQ). The participants were not told about any patterns in the task. In SMP, stimuli were presented in a simple order of button presses 4-3-2-1-4-3-2-1. In RND, stimuli were presented in pseudorandom order, preventing learning in this condition. Pseudorandom orders were generated using Matlab (The MathWorks, Natick, MA, USA), such that items were not repeated directly after one another. In SEQ, stimuli were organized in an 8-items-sequence in a second order predictive manner, which means that pairs of consecutive stimuli, rather than single stimuli (which would be first-order predictive) are not followed by some other stimuli, thereby preventing learning by pairwise associations (Curran, 1997). The task contained a total number of 11 blocks. These comprised of one SMP block at the beginning or the end of the task (counter-balanced between subjects and sessions), and 5 RND blocks and 5 SEQ blocks presented in alternating order (see Fig. 1D). Each of the SEQ blocks contained 15 repetitions of the 8-element sequence summing to a total of 120 trials per block. The SMP block contained 10 repetitions of the simple sequence summing to a total of 80 trials. Each of the RND blocks contained 80 trials. A 20 s break was introduced between the blocks during which participants were instructed to fixate on a black cross in the center of the screen and to try not to move.

### 2.4. Completion task

In this task, the same stimuli were presented as in the SRTT (see above), with the exception that only sequence trials were shown. The 8-element sequence was repeated 32 times. In each repetition, one of eight regular trials, was substituted by a completion trial. In a completion trial, no target square was shown and instead a question mark was displayed above four black squares. Subjects had to press the corresponding button to the square, which they believed would follow next in the sequence. Each position in the sequence was tested four times, producing 32 completion trials. After guessing, subjects were asked whether they were sure of their choice and had to give a YES/NO answer. We were thus able to differentiate between a correct response and a correct assured response.

### 2.5. Transcranial direct current stimulation protocol

Anodal transcranial direct current stimulation (tDCS) was delivered through two 3 cm × 3 cm MR-compatible rectangular rubber electrodes, 3 mm thick. The electrodes were covered with 2 mm of Ten20® conductive paste (Weaver and company, Aurora, CO, USA). The paste was evenly distributed across the electrode surface using a specialized custom-made device. The electrodes were connected through MR-compatible cables to an inner filter box lying inside the MRI. The cables contained two 5 kΩ resistors to reduce induction voltage due to high RF impulses. The inner box was then connected via LAN cable to an outer filter box, placed outside of the scanner room in close proximity to the cable exchange pipe between the MR control room and the scanner room. The outer filter box absorbed RF noise mostly between 50 and 140 MHz which can be produced by the scanner resonance frequency. The outer filter box was then connected to the Neuroconn DC-stimulator PLUS device (neuroCare Group GmbH, Ilmenau, Germany). Impedance measured directly between stimulator and electrodes was kept below 3 kΩ (outside of scanner). Prior to electrode placement, the skin surface was treated with Nuprep (Weaver and company, Aurora, CO, USA), a mild abrasive gel for lowering skin impedance. Electrodes were placed either to stimulate right cerebellum (rCB) or left M1 (lM1) based on computational modelling of the optimal placement of stimulation electrodes (supplementary materials, section 1). For rCB tDCS, the anodal electrode was centred 3 cm right lateral to the inion, and the cathodal electrode was positioned on the right mandibula (Fig. 1B). For lM1 tDCS, the anodal electrode was centered at FC3 and the cathodal electrode at CP3 according to the 10-20 EEG system, creating the maximum current density over C3 (Fig. 1B). For sham, electrodes were placed either with lM1 or rCB montage. The electrodes were fixed on the head using rubber-straps. The current was set for both lM1 and rCB tDCS to 1 mA resulting in a current density of 0.11 mA/cm^2^, which is in the range of standard 2 mA cerebellar tDCS with larger electrode surface (< 0.08 mA/cm^2^) (Ferrucci et al., 2015).

TDCS initiated with the first trial in the task. For lM1 and rCB tDCS, the current was ramped up for 30 s, remained at 1 mA for 20 min, and ramped down for 30 s. Note that duration of the SRTT varied with subjects’ individual task performance. Thus, sometimes stimulation resumed before task completion and rarely after task completion. However, the non-overlapping time between SRTT and tDCS was never longer than 2 min. For sham, the current was ramped up for 30 s, remained at 1 mA for 60 s, and ramped down for 30 s.

### 2.6. Behavioral analysis

One subject was excluded from the behavioral analysis due to a malfunction in the keypad in one of the experimental sessions, resulting in a sample of 24 subjects for this analysis. We computed reaction times (RTs) and error-rates for each of the experimental conditions. Both wrong button-presses and missing responses were regarded as Errors. RTs were averaged across mini-blocks of 40 trials, corresponding to five repetitions of the 8-element sequence in SMP and SEQ. This sub-division resulted in two mini-blocks for each RND and SMP block and three mini-blocks for each SEQ block (see Fig. 1D). For RT analysis, we excluded trials in which the participants pressed a wrong button as well as trials in which RTs were either longer than 1000 ms or deviated by more than 2.7 standard deviations (SD) from their average response time (corresponding to p < 0.01). To specifically measure motor sequence learning, we measured the RT difference between SEQ blocks, in which sequence learning occurs and RND blocks in which no sequence learning is expected due to pseudorandom presentation of visual stimuli. Importantly, if subjects had learned the underlying SEQ, they would show a significant speeding up when re-exposed to the SEQ. This speeding up should increase with time. We defined three different time windows for the analysis: Early time window, Mid time window, and Late time window (see Fig. 1D). We looked specifically for the first transfer from RND to SEQ in each of these time windows, and entered averaged normally distributed RTs into a 3 × 2 × 3 repeated measures ANOVA (rmANOVA) with factors STIM (rCB, lM1, sham), COND (SEQ, RND) and TIME (Early, Mid, Late). Note that COND x TIME interaction reflects the progress of sequence learning with time and the effect of tDCS on this progress can be assessed by testing STIM x COND x TIME interaction.

The error-rate of each task block (SEQ and RND) was computed as the number of Errors divided by the number of trials in each block (RND – 80, SEQ – 120). As error rates were not distributed normally, we tested for learning in each stimulation protocol by comparing error rates in RND to SEQ, averaged across all time windows, using the Wilcoxon signed rank test.

Analysis of the completion task was performed by assessing the differences between percentage correct and percentage correct assured responses between stimulation protocols using the Wilcoxon signed rank test.

### 2.7. MRI data acquisition

During stimulation and SRTT performance, participants were lying supine in a magnetic resonance imaging (MRI) scanner with their right arm comfortably placed on a padded armrest and the right hand placed on a MR-compatible keypad. The keypad was fixed to a strap wrapped around their upper legs. To prevent head motion, the head was fixed with foam pads between the ears and the head coil. The foam pads also provided subjects with noise protection. The experimental stimuli were presented using Presentation® (NeuroBehavioral Systems) software on a 32” LCD monitor (NordicNeuroLab) with a resolution of 1920 × 1080 pixels and an active area of 698.4 mm × 392.9 mm, located at the back of the MRI scanner and visible through a tilted mirror integrated into the head coil.

The MRI data were recorded using a 3T Siemens MAGNETOM Skyra head-scanner at the Center for Brain Behavior and Metabolism, University of Lübeck. A high resolution T1-weighted gradient-echo structural image was acquired (image matrix: 256 × 256; 192 sagittal slices of 1mm thickness; TR = 1900 ms; TE = 2 ms). Functional MRI data (T2*) were collected using blood oxygen level dependent (BOLD) contrast. During stimulation and task performance, a gradient-echo EPI sequence was used with the following specifications: repetition time TR = 2000 ms, echo time TE = 25 ms, flip angle = 90°, matrix size 64 × 64, FOV = 192 × 192 mm with a whole brain coverage, 40 axial ascending slices of 3 mm thickness and 25% gap and in-plane resolution of 3 mm × 3 mm, iPAT factor of 2.

### 2.8. FMRI pre-processing and statistical analyses

Preprocessing of fMRI data was done using the SPM12 software package (http://www.fil.ion.ucl.ac.uk/spm/) and comprised slice timing correction, realignment to correct for head motion artifacts, co-registration to T1 structural image, segmentation, normalization to Montreal Neurological Institute (MNI) template brain image, smoothing with a Gaussian kernel of 8 mm full width half maximum and resampling of functional images to 3 × 3 × 3 mm. Imaging data was subsequently modeled using the general linear model (GLM) in a block design manner. Analyses were performed for each of the stimulation protocols (lM1 tDCS, rCB tDCS, sham) and each subject. First level GLM analysis thus contained 11 experimental blocks (5 × SEQ, 5 × RND and 1 × SMP) modeled as a box function with the duration of each block, and convolved with a hemodynamic response function. Movement related parameters from the realignment process were included in the GLM as regressors of no-interest to account for variance caused by head motion. We applied a high-pass filter (256 s) to remove low-frequency noise. First-level contrast images were generated using a one-sample t-test (against baseline) in three different time windows: Early, Mid and Late (Fig. 1D). were analyzed on the second level using a flexible factorial design accounting for effects of stimulation (STIM: lM1 tDCS, rCB tDCS and sham) and condition (COND: SEQ and RND) as well as interactions. Note that COND x STIM interactions in each time window reflects the specific effect of tDCS on sequence learning. We therefore explore learning-related changes in activity due to stimulation, defined as the differences in the BOLD signal between SEQ and RND blocks. Statistical significance was established by thresholding the statistical maps at a whole-brain uncorrected voxel-level threshold of p = 0.001, followed by cluster-level correction of p < 0.05, family-wise error corrected for the entire cluster. In addition, we used rfxplot (Gläscher, 2009) to extract signals from clusters showing significant activity, using 8 mm spheres around the peak activation.

In order to test whether differences in the BOLD signal between conditions were associated with individual learning ability in each of the stimulation protocols (lM1 tDCS, rCB tDCS and sham), we performed a regression analysis for each time window (Early, Mid, Late) separately, between the averaged RT difference (SEQ-RND) and a contrast image for the condition differences, in each subject and each session. The resulting statistical maps thus represent correlations between subject specific performance and changes in learning-related (SEQ-RND) activity. Similar to the analysis above, significance was an uncorrected voxel-level threshold of p = 0.001, and family-wise error cluster-level correction of p < 0.05.

Next, we tested whether correlations between regional activity and performance under lM1/rCB tDCS significantly differed from similar correlations under sham. First, we extracted activity under sham from the same location and the same time window showing significant correlation under real tDCS. Then, we calculated the correlation between this activity and performance during sham. Finally, we Fisher transformed the correlation coefficients *r* for real tDCS (*r*_*1*_) and sham (*r*_*2*_) as follows: 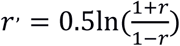, and computed the z statistic: 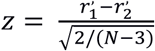, where N = 24 (number of subject).

### 2.9. Dynamic causal modelling and model space specifications

We used dynamic causal modeling (DCM, Friston et al., 2003) as implemented in SPM12 (v. 7219), version DCM12, to investigate the influence of tDCS on effective connectivity within the cortico-striato-cerebellar network. DCM is useful to describe directional interactions between brain regions underlying a specific experimental manipulation. Dynamic changes in regional activity are thus described using a one-state bilinear system of differential equations:

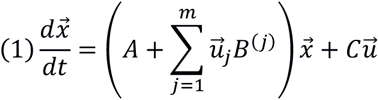

Where 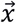 represents a neuronal state vector that changes over time (dt) and 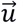 an input vector. We constructed the input vector 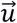 as a stick function of single events corresponding to the onset of each trial. 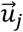 is the j-th experimental input to the modelled system. Our models had only one input such that j = m = 1. *A* represents the endogenous (context independent) interactions within the network, *B* represents the modulatory (context dependent) influence on connections by input/context 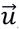, and *C* the extrinsic effects of input 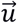 on activity. *A, B, C* parameters are rates of change (rate constants) which are estimated for a complete time series. If A is positive, then the connection is excitatory, e.g. region 1 increases activity in region 2 with time. If A is negative, then the connection is inhibitory. Similarly, B is the increase or decrease in connectivity from region 1 to region 2 with time, due to the experimental condition j.

Note that the input represents stimulus-response associations that take place in each trial. Therefore, the cerebellum was defined as the input node based on its importance for proprioception (Bhanpuri et al., 2013). In previous fMRI studies of motor sequence learning, we found that right cerebellum as input node leads to highest exceedance probability (Tzvi et al., 2017, 2015, 2014) and therefore chose this node as input here as well. It is also likely that the driving input is mediated by un-modelled regions such as the visual cortex.

We specified 60 different models (supp. Fig. 2) based on a-priori assumptions from previous DCM studies of the motor system (Grefkes et al., 2008; Pool et al., 2013), our studies of motor learning networks (Tzvi et al., 2020, 2017, 2015, 2014)as well as theoretical predictions of motor learning (Doyon et al., 2009; Hikosaka et al., 2002) stating that the early learning phase should activate a cortico-striatal-cerebellar network in MSL. These 5-node network models contained left motor cortical areas: primary motor cortex (M1), supplementary motor area (SMA) and premotor cortex (PMC) as well as left putamen and right cerebellum. All VOIs were fully connected. Note that the input and connections specified here do not necessarily represent anatomical input and connectivity but rather a “net effect”.

**Figure 2.**
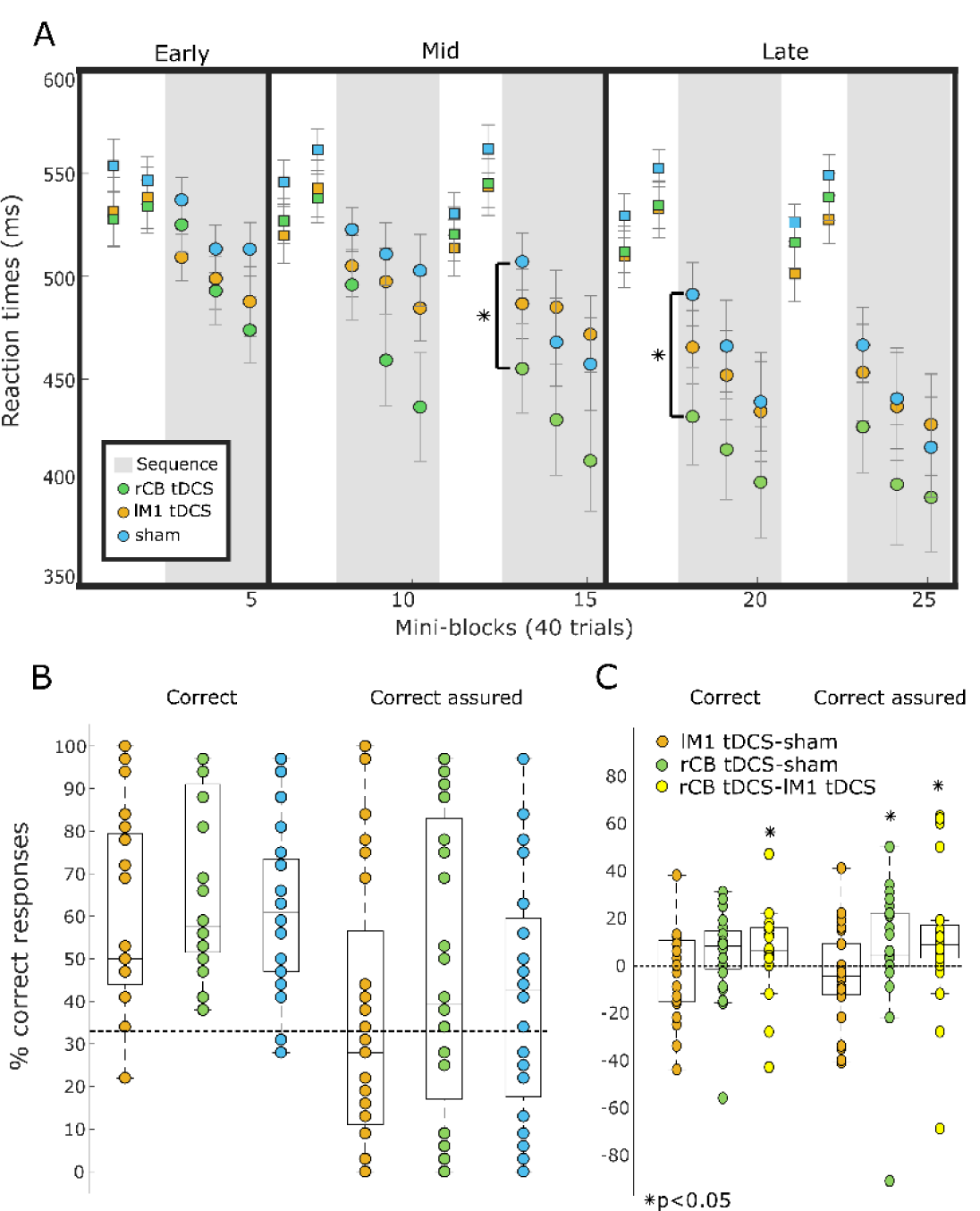
Behavioral results. **A** Reactions times aggregated across subjects and sessions. Each point in the graphic represents average data of 40 trials (mini-block). Error bars are standard error of the mean across subjects. Sequence blocks are marked with a grey shade. Significant differences are marked with a star. **B** Boxplots for percentage correct responses in the completion task. Individual subjects are depicted with a filled circle. Chance level is shown with a dashed black line. Significant differences between protocols are marked with a star (posthoc t-tests, p < 0.05).

We previously found that interactions between M1 and PMC to cerebellum, as well as putamen-cerebellar interactions were negatively modulated by motor sequence learning. We therefore hypothesized that these connections would be modulated by learning and by tDCS to lM1 or rCB. Note that we chose to include only left hemispheric (and right cerebellar) nodes as subjects performed the task with the right hand and in order to limit the model space which exponentially grows the more nodes are included (and thus the computational effort). A separate analysis of a 4-node network comprising bilateral M1 and cerebellum showed very poor fit (supplemental materials, section 7).

For each model, the differential system of equations described in (1), was inverted and together with a biophysically motivated hemodynamic model, an estimated BOLD signal was produced. This modelled BOLD signal was then iteratively fitted to the real data through a gradient ascent on the free-energy bound.

We describe the results of the DCM analysis on two levels: on a model level and on a parametric level. First, for each of the stimulation protocols (rCB tDCS, lM1 tDCS and sham), we selected a “winning” model out of a candidate set of equally plausible models based on its protected exceeding probability using Bayesian model selection. Second, we used Bayesian model averaging (BMA) to compare connectivity parameters between rCB tDCS or lM1 tDCS and sham using random-effects analysis across subjects. Note that BMA produces a posterior distribution over connection strength across all models which is used to compute the average connection strength common to all subjects.

#### 2.9.1 Time series extraction

We specified left primary motor cortex (M1), left putamen (Pu), left supplementary motor area (SMA), left premotor cortex (PMC) and right cerebellum (CB) as volumes of interest (VOIs) similar to our previous work (Tzvi et al., 2017, 2015). Time series were extracted from significant voxels (p < 0.05, uncorrected) in the task vs. baseline individual contrasts, across all experimental blocks and stimulation sessions, in order to account for both learning-and non-learning related changes in the BOLD signal. The coordinates of the sphere center for each VOI were selected based on the local maxima of the group level task vs. baseline contrast in each stimulation session (see supp. Table 1). For each individual subject, the sphere center of each VOI was moved to the closest suprathreshold voxel, which was always kept within a Euclidean distance 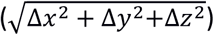 of 10 mm from the original sphere center. Using the xjview toolbox (http://www.alivelearn.net/xjview) and AAL brain atlas, we verified that sphere centers for all subjects were within the regions of interest. For M1, sphere centers were kept within BA4 and precentral gyrus. For PMC, sphere centers were kept within BA6 and middle frontal gyrus. For cerebellum, sphere centers were kept within lobules IV-V. In four subjects, the coordinates for left putamen were further than 10 mm from the original sphere center, but were still within putamen. These subjects were nevertheless included in the DCM analysis for statistical power. In one subject, we couldn’t find any activations in the specified regions of interest in one of the sessions. This subject was therefore excluded from the DCM analyses resulting in a sample of 23 subjects.

**Table 1.** Regional activations: tDCS vs. sham in Mid time-window (p < 0.05, cluster-level family-wise error corrected)

| Region | MNI-coordinates |  |  | t-value | n-voxels |
| --- | --- | --- | --- | --- | --- |
| L inferior frontal gyrus | -34 | 30 | 18 | 5.24 | 66 |
| R inferior parietal lobule | 48 | -34 | 52 | 4.95 | 37 |
| L cerebellum lobule VI | -24 | -58 | -34 | 4.81 | 43 |
| L cerebellar crus I | -38 | -58 | -34 | 4.46 | 71 |
| R primary motor cortex | 48 | -30 | 60 | 4.25 | 67 |

Using a singular value decomposition procedure implemented in SPM12, we computed the first eigenvariate across all suprathreshold voxels within 6 mm radius from the sphere center for each subject. We chose a relatively large radius for the spheres to avoid including signals in the DCM analysis which are derived from VOIs with very few voxels and thus are more susceptible to noise. Time series were then detrended and sharp improbable temporal artifacts were smoothed by an iterative procedure implementing a 6-point cubic-spline interpolation. Finally, we estimated the explained variance of the signals we extracted from each of the VOIs by computing the proportion of the first eigenvariate in the signal. In all subjects and all VOIs, signals had more than 40% variance explained, which means that the first eigenvariate explained much of the signal variance in the different VOIs. Note that variance in lM1 and rCB VOIs did not differ between stimulation protocols or in comparison to sham (p > 0.1), suggesting that SNR in regions close to the stimulation electrodes was not affected by the stimulation.

#### 2.9.2 Analysis of modulatory parameters using BMA

In each subject and each session, we estimated the averaged posterior distribution of modulatory effects across all models using BMA. We tested for the effect of tDCS on modulatory parameters during learning using 2 × 2 rmANOVA with factors STIM (rCB tDCS or lM1 tDCS vs. sham) and COND (SEQ, RND) in specific time windows. In addition, we analyzed correlations between modulatory parameters and RT differences, as well as percentage accuracy as a measure for learning in each stimulation protocol (lM1 tDCS, rCB tDCS, sham).

## 3. Results

Subjects tolerated the tDCS protocol well without any adverse effects. Blinding through sham tDCS, assessed with questionnaires, was successful. Please refer to supplementary materials, section 2, for further details.

### 3.1. Right cerebellar tDCS enhances learning in the serial reaction time task

To assess changes in motor sequence learning due to rCB and lM1 tDCS, we subjected the reaction times (RTs) of mini-blocks at the first transfer from RND to SEQ (see methods for details), to a repeated measures ANOVA (rmANOVA) with factors STIM (rCB, lM1, sham), COND (SEQ, RND) and TIME (Early, Mid, Late). We found a main effect of COND (F_1,23_ = 77.5, p < 0.0001), a main effect of TIME (F_2,46_ = 22.7, p < 0.0001) as well as COND x TIME interaction (F_2,46_ = 22.2, p < 0.0001), demonstrating that learning as measured by RT difference between SEQ and RND, increased with time in all experimental sessions (see Fig. 2A). Importantly, we found a COND x STIM x TIME interaction (F_4,92_ = 3.2, p = 0.02) revealing specific effects of stimulation on differences in RT between conditions. A separate 2 × 2 × 3 rmANOVA comparing rCB to sham revealed a COND x STIM x TIME interaction (F_2,46_ = 3.3, p < 0.05) as well, with no such interaction evident in a similar rmANOVA comparing M1 to sham (p > 0.2). Interestingly, a rmANOVA comparing rCB tDCS to lM1 tDCS led to a significant COND x STIM x TIME interaction (F_2,46_ = 3.8, p = 0.03), suggesting that lM1 and rCB tDCS have different effects on learning.

To specifically test at which time point performance differed most between the stimulation protocols, we compared RTs of the first mini-block of each SEQ block across stimulation sessions (lM1 tDCS vs. sham, rCB tDCS vs. sham and lM1 tDCS vs. rCB tDCS). We expected to find differences due to stimulation in the Mid and Late time windows. These tests revealed that participants were significantly faster during SEQ performance following rCB tDCS (compared to sham) in the second transfer at Mid time window (t_23_ = 2.1, p = 0.04, Fig. 2A) and the first transfer at Late time window (t_23_ = 2.2, p = 0.03, Fig. 2A). No differences were found between M1 tDCS and sham (all p > 0.1), as well as between CB and M1 tDCS (all p > 0.2). Note that these tests were not corrected for multiple comparisons. These results suggest that cerebellar tDCS led to faster learning in Mid and Late time windows, while lM1 tDCS had no effect on learning.

We found no difference in initial task performance, measured as RT average in the first block, between the stimulation sessions (lM1 tDCS vs. sham, rCB tDCS vs. sham and lM1 tDCS vs. rCB tDCS: all p > 0.2).

As subjects performed the SRTT a total of three times in three sessions, we also tested whether sequence learning significantly improved across sessions (see detailed analysis in the supplementary materials, section 4). A significant COND x TIME x SES (session order) interaction (p = 0.002) suggested that indeed learning improved across sessions. A post-hoc analysis in each of the sessions separately, testing for stimulation effects using a mixed model revealed a COND x TIME x STIM interaction in the last session (F_4,42_ = 2.9, p = 0.04), suggesting that the observed improved learning under rCB tDCS is mainly driven by better performance in the last session of the SRTT.

As error rates were not distributed normally, we were not able to submit them to a measures rmANOVA similarly to the RTs. We tested for learning in each stimulation protocol (M1 tDCS vs. sham; rCB tDCS vs. sham) by comparing error rates in RND to SEQ, averaged across all time windows, using the Wilcoxon signed rank test. We found larger error rates in RND compared to SEQ (M1: p = 0.007, CB: p < 0.001, sham: p = 0.003) across all stimulation protocols. No differences were observed when comparing between stimulation protocols (all p > 0.1).

### 3.2. Right cerebellar tDCS enhances performance in the completion task

An additional subject was excluded from this analysis due to a technical error resulting in a sample of 23 subjects. Note that this subject was not excluded from the following fMRI analysis during SRTT performance. Subjects were tested on 32 completion trials. Figure 2B shows the distribution of correct and correct assured (a correct response followed by a “yes” answer to the question: “are you sure?”) responses across subjects and stimulation protocols.

First, we examined in each session, how many subjects were better than chance level of 33% using the Wilcoxon signed rank test as data were not distributed normally (see single data points in Fig. 2B). Across all stimulation protocols subjects were significantly better than chance (p < 0.001): for lM1 tDCS, 23 out of 24 (96%) were better, for rCB tDCS 24 out of 24 (100%) were better and for sham, 21 out of 24 (88%) were better than chance when guessing the next button press. We then tested how many subjects had correct assured responses that were higher than chance level to identify if subjects have gained some explicit awareness of the sequence. Here, the group had mixed responses: following lM1 tDCS 10 out of 24 subjects (42%) were above chance level, for rCB tDCS and sham 15 out of 24 were above chance level (63%).

We then directly tested differences between percentage correct and percentage correct assured responses between stimulation protocols using the Wilcoxon signed rank test (lM1 tDCS vs. sham, rCB tDCS vs. sham and lM1 tDCS vs. rCB tDCS). The results are presented in Figure 2B and 2C (significant differences are marked with a star). Following rCB tDCS, subjects were significantly better compared to lM1 tDCS (Z = 2.20, p = 0.03), and marginally better than sham (Z = 1.74, W = 180, p = 0.08). There were no differences between lM1 tDCS and sham in terms of correct responses (Z = -0.6, p = 0.55). In terms of correct assured responses, following rCB tDCS subjects were significantly better compared to lM1 tDCS (Z = 2.52, p = 0.01) and compared to sham (Z = 2.21, p = 0.03). There were no differences between lM1 tDCS and sham in terms of correct assured responses (Z = 1.0, p = 0.31). Together these results show that subjects were better in securely and correctly guessing the next button press under rCB tDCS compared to both lM1 tDCS and sham.

### 3.3. Imaging results cerebellar tDCS: learning-related activity increases during rCB tDCS

To investigate changes in brain activity resulting from rCB tDCS at different time windows (Early, Mid, Late), we performed a flexible factorial analysis with factors STIM (rCB tDCS vs. sham) and COND (SEQ, RND) in each time window (Early, Mid and Late) as the effect of STIM may differ with time. First, we report the main effect of STIM, which reflects general task-related changes in activity resulting from rCB tDCS. There were no significant changes in activity comparing rCB tDCS to sham on a corrected cluster level (see methods, section 2.8). In Early, a tendency for decreased activity in left M1 was evident during rCB tDCS compared to sham (p_peak_ < 0.001, p_cluster_ = 0.02, t = 4.0, N_voxels = 62). In Late, activity tended to increase in right M1 during rCB tDCS compared to sham (p_peak_ < 0.001, p_cluster_ = 0.02, t_peak_ = 4.2, N_voxels = 62).

Next, we investigated learning-related changes in brain activity, i.e. the effect on condition differences (SEQ-RND) due to rCB tDCS, at different time windows. Here, we report STIM x COND interactions of the flexible factorial analysis above, reflecting changes in learning-related activity between rCB tDCS and sham. Please refer to supplementary materials, section 5, for an exploratory analysis comparing rCB tDCS to lM1 tDCS. To better understand these interactions, we used rfxplot to extract signals from these regions, using 8 mm spheres around the peak activation. We then plotted the parameter estimates, representing percentage activity change for each of the time windows in each condition and stimulation protocol.

Significant changes in learning-related (SEQ-RND) activity during rCB tDCS compared to sham were observed only in the Mid time window. These effects were observed in right M1, right inferior parietal lobe (IPL), left inferior frontal gyrus (IFG), left cerebellar crus I, and left cerebellar lobule VI (see Table 1). A cluster in left M1 (p_peak_ < 0.001, p_cluster_ = 0.01, t = 4.0) and left IPL (p_peak_ < 0.001, p_cluster_ = 0.02, t = 4.0) were evident on a trend level. The activation maps in Figure 3A show this COND x STIM interaction in left cerebellar lobule VI, left inferior frontal gyrus (IFG), and right M1. In Figure 3B the percentage signal change extracted from these regions is displayed across all time windows.

**Figure 3.**
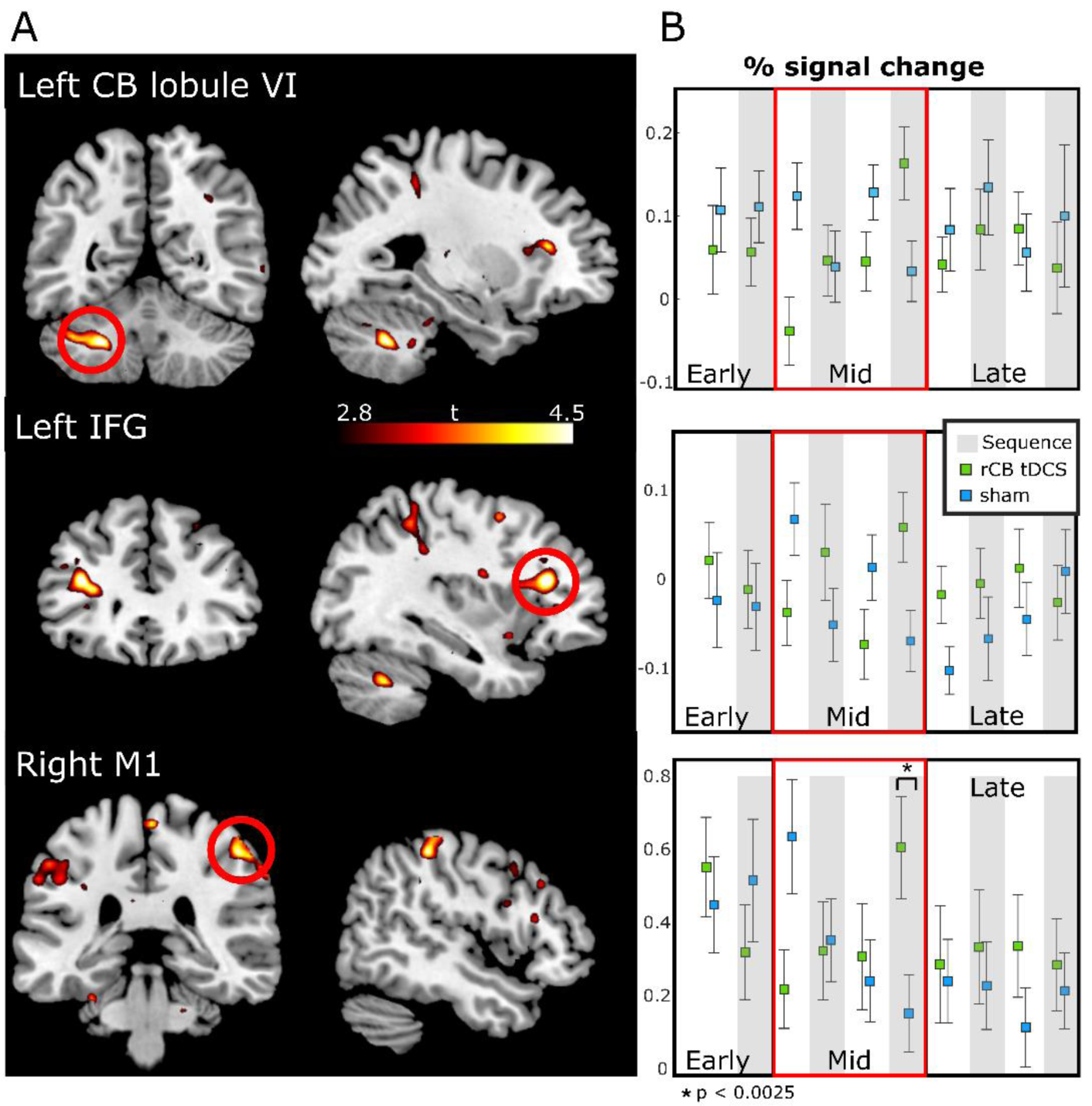
Effects of rCB tDCS on learning-related activity in Mid time window reflected by COND (SEQ, RND) x STIM (rCB tDCS, sham) interactions. **A** Activation maps shown on a p < 0.001 voxel-level threshold with p < 0.05, family-wise error corrected on the cluster level for multiple comparisons. **B** percentage signal change in each condition and stimulation extracted from the activations marked in A with a red circle. Note that activations were evident in Mid, Early and Late time windows are presented for reference. Asterisk denotes significant difference in post-hoc t-tests (see Table 2).

Visual inspection suggested that the interaction was driven by larger activity in sham compared to rCB tDCS during RND, and larger activity in rCB tDCS compared to sham during SEQ. Note that there were no significant activations for the reverse contrast (the opposite effect). We therefore conducted post-hoc t-tests comparing the percentage signal change in rCB tDCS to sham, for each of the five regional activation clusters (correcting for multiple comparisons, see Table 2). We found that activity in rM1 was larger in rCB tDCS compared to sham in the second SEQ block (see Fig. 3B). In the other regions, activity tended (p < 0.05, uncorrected) to be larger in rCB tDCS compared to sham in the second SEQ block as well. During the first RND block on the other hand, activity in right M1, IPL, and left cerebellar crus I and lobule VI tended (p < 0.05, uncorrected) to be larger during sham compared to rCB tDCS (see Fig. 3B). Correlation analysis between subject-specific performance and learning-related (SEQ-RND) activity changes revealed no clusters for rCB tDCS.

**Table 2.** Regional activations: post-hoc t-tests in Mid time-window

| Region | rCB tDCS > sham |  |  |  |  |  |  |  |
| --- | --- | --- | --- | --- | --- | --- | --- | --- |
|  | SEQ1 |  | SEQ2 |  | RND1 |  | RND2 |  |
|  | t | p | t | p | t | p | t | p |
| R M1 | -0.2 | 0.8 | 4.0 | 0.0006** | -3.0 | 0.006* | 0.6 | 0.5 |
| L IFG | 1.5 | 0.15 | 2.3 | 0.03* | -1.9 | 0.07 | -2.2 | 0.04* |
| L CB lb VI | 0.14 | 0.9 | 3.1 | 0.005* | -3.1 | 0.005* | -1.8 | 0.09 |
| R IPL | -0.1 | 0.9 | 3.1 | 0.005* | -3.0 | 0.007* | 0.09 | 0.9 |
| L CB cr I | 0.6 | 0.6 | 2.2 | 0.04* | -3.0 | 0.007* | -1.4 | 0.2 |

\*\* $p < 0.05$ , Bonferroni corrected for 2 Conditions x 2 blocks x 5 regions comparisons: $p < 0.0025$
\* $p < 0.05$ , uncorrected

Summarizing the results of rCB tDCS (compared to sham) on regional activity, we found a general task-related decrease in left M1 during Early, and increase in right M1 during Late (on a trend level). Learning-related (SEQ-RND) activity increased during Mid in right M1, left cerebellum crus I and lobule VI, right IFG, and right IPL.

### 3.4. Imaging results M1 tDCS: enhanced learning is associated with increased activity

Despite no evidence for a behavioral effect of lM1 tDCS on learning, we performed an exploratory analysis comparing lM1 tDCS to sham using the flexible factorial analysis described above, with factors STIM (lM1 tDCS vs. sham) and COND (SEQ, RND) in each time window (Early, Mid and Late). For the main effect of STIM, we found no significant changes in activity comparing lM1 tDCS to sham. During Mid, activity tended to increase in right cerebellar crus II during lM1 tDCS compared to sham (p_peak_ < 0.001, p_cluster_ = 0.01, t_peak_ = 5.2, N_voxels = 72). There were no STIM x COND interactions.

Next, we analyzed possible correlations between subject-specific performance, reflected by RT differences between conditions and learning-related (SEQ-RND) changes in activity during lM1 tDCS. We found a rather weak association between decreased activity in left middle occipital cortex and better learning during Early (R = 0.22, see Table 3) and a strong association between decreased activity in right paracentral lobule with better learning during Mid (R = 0.93, see Table 3). In Late, we found that activity increase in right M1, right SMA and bilateral middle frontal gyrus was associated with better learning (Fig. 4A and Table 3). This means that subjects who performed very well during lM1 tDCS had also stronger activations in these regions during sequence performance (compared to random), whereas subjects with worse performance during lM1 tDCS had weaker or no activations in these regions (see Fig. 4B). Note that the correlations presented in Figure 4A-B remain significant (all p < 0.0001) also when removing the three extreme behavioral points (see Fig. 4B). A similar correlation analysis in sham showed that activity decrease in left temporal cortex and right inferior occipital cortex was rather weakly associated (R = -0.22, -0.03 resp., see Table 3) with better learning in the Early time window. We further tested whether the correlation coefficients between lM1 tDCS and performance differed from the correlation coefficients between sham and performance in the regions above, during the respective time-windows, using the Fisher-transform. Correlation coefficients were significantly different in all regions, except for right SMA and left middle frontal gyrus during Late (see Table 3).

**Table 3.**
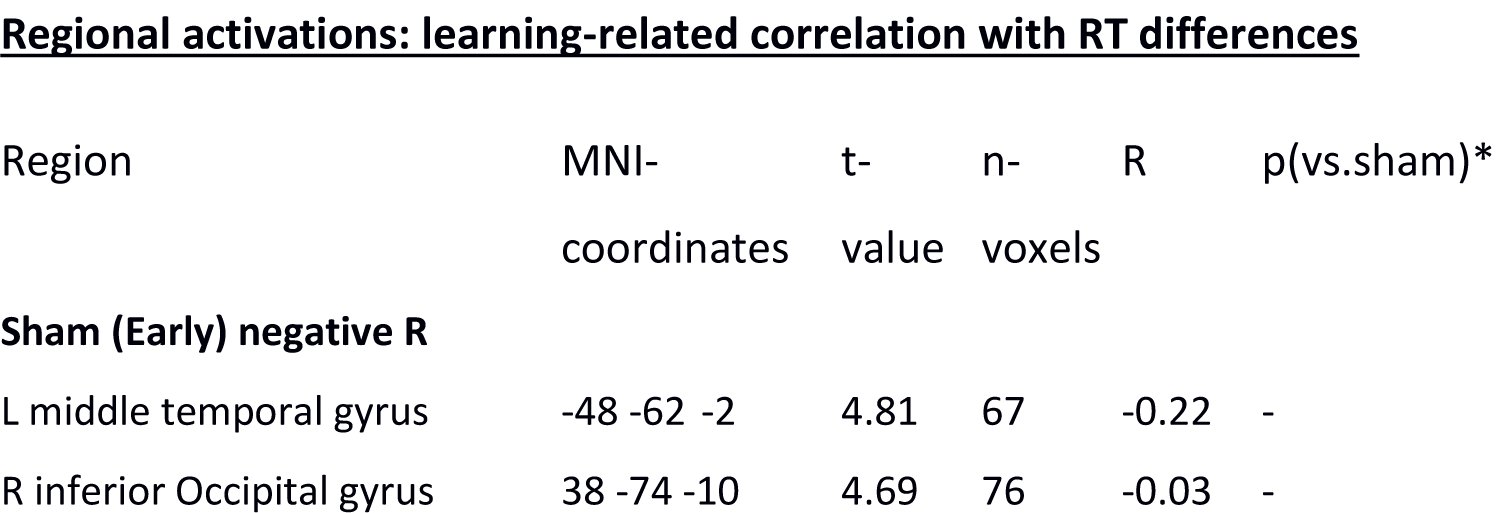
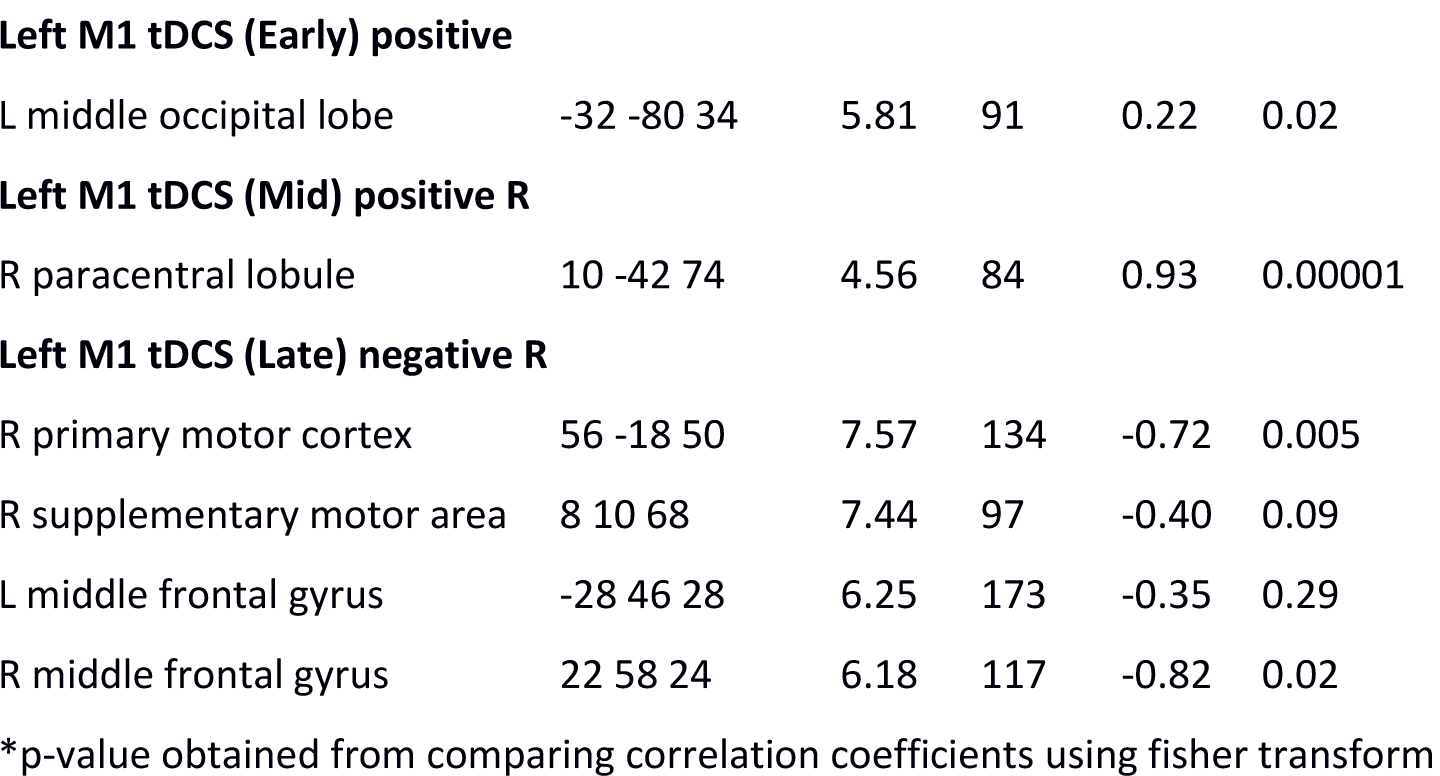
Regional activations: learning-related correlation with RT differences

**Figure 4.**
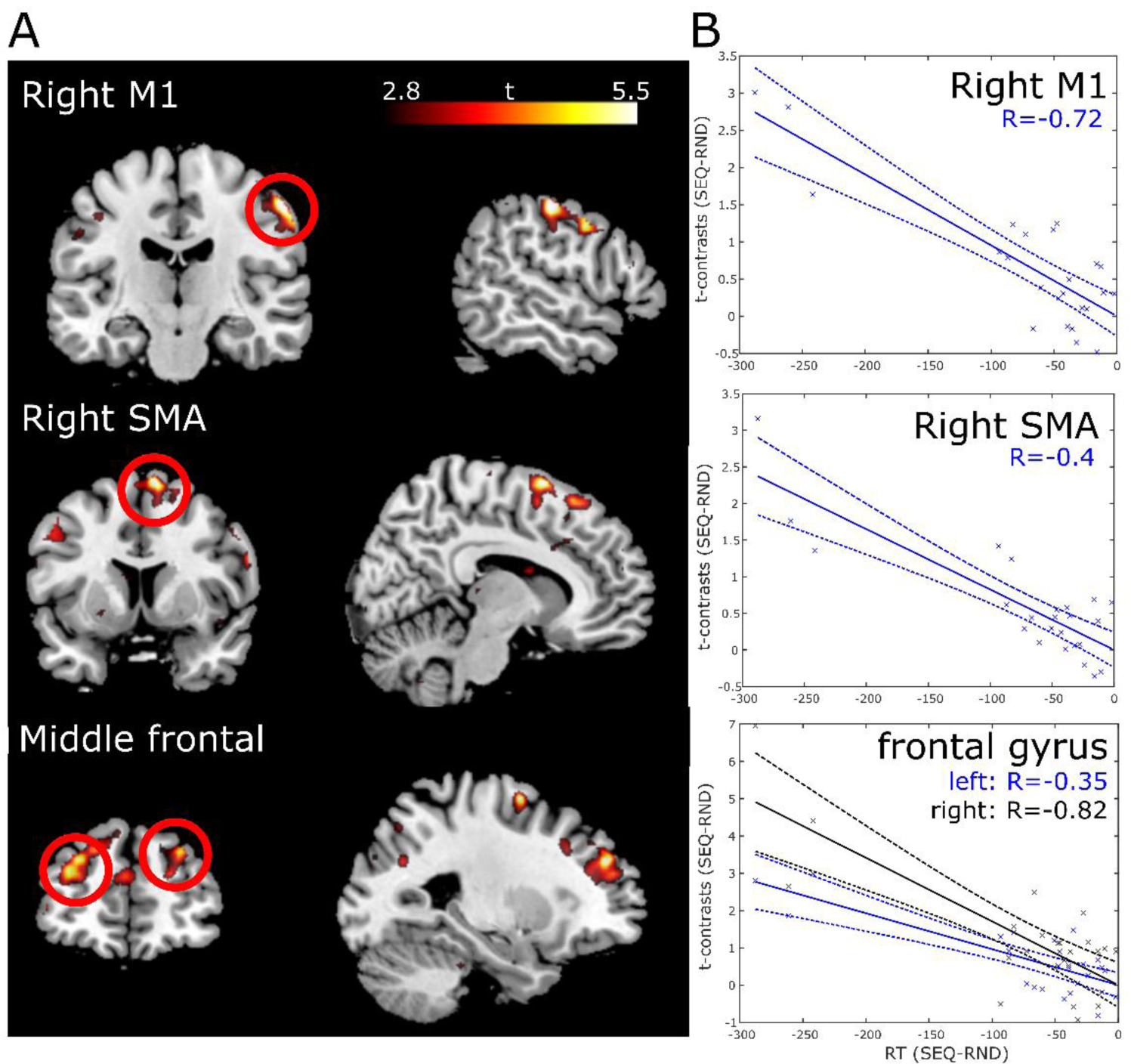
Better learning correlates with increased activity during lM1 tDCS in Late time window. **A** activations reflecting negative correlations between RT (reaction times) differences and individual t-contrasts for learning-related (SEQ-RND) activity in Late under lM1 tDCS. Activations are shown on a p < 0.001 voxel-level threshold with p < 0.05, family-wise error corrected on the cluster level for multiple comparisons. T-values represent the strength of the correlation. **B** correlations between averaged t-contrasts for the cluster presented in A and RT differences in Late under lM1 tDCS.

Summarizing the results of lM1 tDCS (compared to sham) on regional activity, we found, on a trend level, a general task-related increase in right cerebellar crus II during Mid. No learning-related (SEQ-RND) changes due to lM1 tDCS were found, as mirrored by absence of behavioral effects (see above). A correlation analysis revealed however, that lM1 tDCS led to increased learning-related (SEQ-RND) activity in right M1, SMA and bilateral middle frontal gyrus, depending on individual learning performance.

### 3.5. DCM results: the optimal cortico-striato-cerebellar model in tDCS and sham

We used DCM to analyze changes in effective connectivity within the cortico-striato-cerebellar network resulting from motor learning and tDCS. To this end, we defined 60 models that differed in modulatory effects on connections between M1, PMC, Pu and CB (supp. Fig. 2), and used the CB VOI as input. Bayesian model selection procedure with a random-effects model was performed for each of the stimulation protocols separately (lM1 tDCS, rCB tDCS and sham), in order to account for changes in network architecture due to tDCS. Figure 5A shows the “winning” model for each stimulation protocol and the comparisons between all 60 models are presented in Figure 5B. For rCB tDCS, the optimal model was model 42, which had modulation of M1, PMC and Pu to CB as well as CB to Pu modulation (Protected exceedance probability: p_ex = 0.22; Bayes omnibus risk for equal model frequencies: BOR = 0.045). For lM1 tDCS, the optimal model was model 2 which had modulation of M1, PMC and Pu to CB (p_ex = 0.23; BOR = 0.048). Lastly, for sham, model 11 which allowed modulation of CB to M1, and Pu to CB connections, had the largest exceedance probability (p_ex = 0.15; BOR = 0.214). Notably, both lM1 and rCB tDCS led to modulation of PMC -> CB and M1 -> CB connection which was lacking in sham. All models had negative modulation of Pu -> CB connection. In supplementary materials, section 6, we provide an analysis of endogenous connections (matrix A) and in supplementary Table 4, a summary of the task input effects to cerebellum (matrix C) across time windows and stimulation protocols.

**Figure 5.**
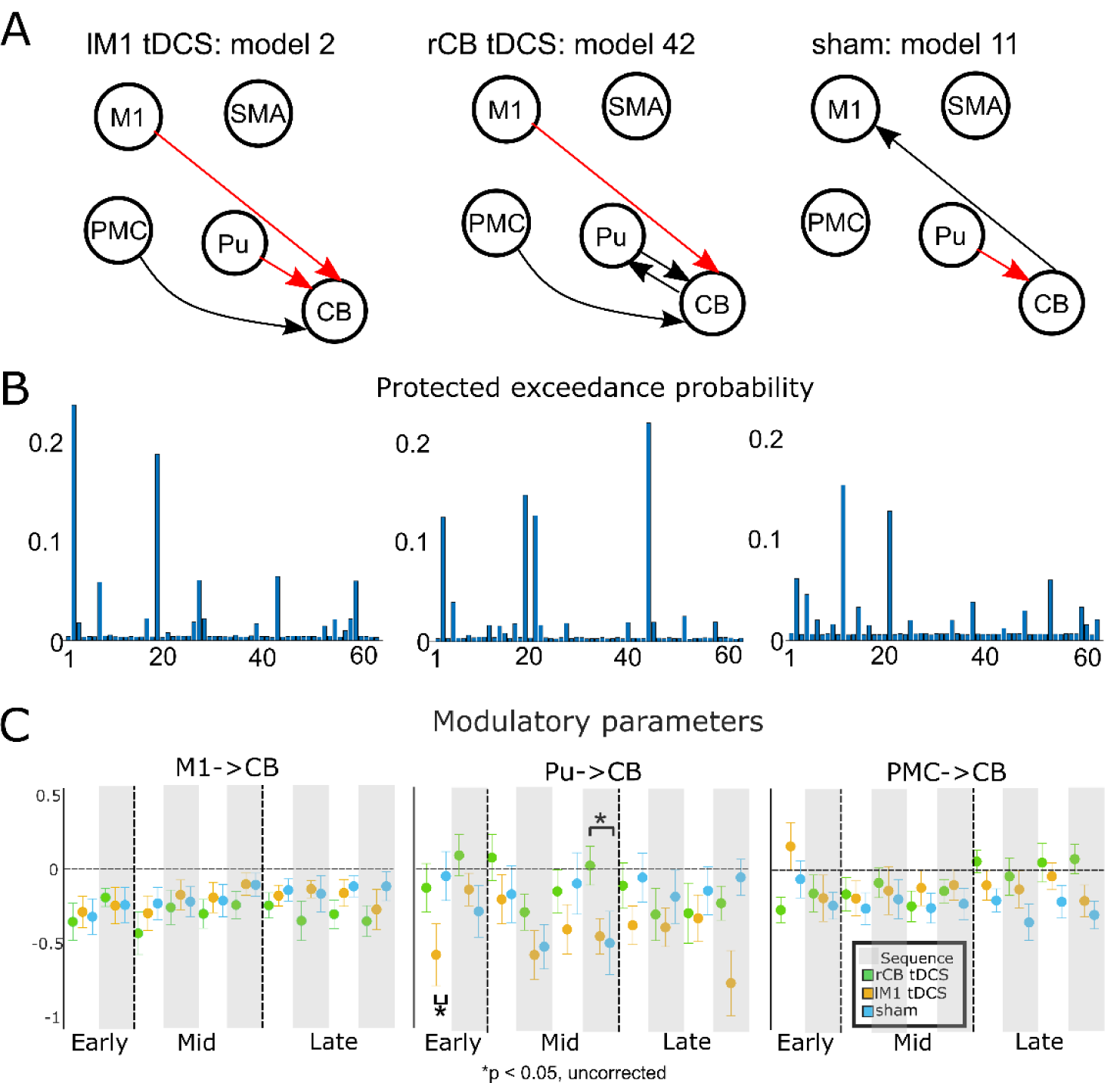
DCM results. **A** The winning models for each of the stimulation protocols showing only modulatory connections. M1 - primary motor cortex, SMA – supplementary motor area, PMC – premotor cortex, Pu – putamen, CB – cerebellum. All VOIs were fully connected. The red arrows mark modulation of connections that were consistently negative. **B** Results of Bayesian model selection across the 60 models (described in supplemental Figure 2) for each of the stimulation protocols. **C** Posterior estimates of modulatory parameters following BMA. COND (SEQ, RND) x STIM (rCB tDCS, sham) interaction was found in the Mid time window for Pu -> CB connection. Post-hoc t-tests show a significant difference, only for SEQ (mark with a grey bar), between rCB tDCS and sham.

### 3.6. Cerebellar tDCS leads to decreased negative modulation in putamen-cerebellar connection

Next, we analyzed the effect of tDCS on modulatory parameters (Fig. 5C). Since each stimulation protocol had a different “winning” model (Fig. 5A), we could not directly compare modulatory parameters between real tDCS and sham, as model structure might influence the posterior estimations. Instead, we used Bayesian model averaging (BMA) to compute a weighted average of parameter estimates across all models. We then proceeded to compare the modulatory effects of Pu -> CB connection across all the stimulation protocols as this connection was presented in all optimal models (Fig. 5A). Note that in Figure 5C we show the averaged modulatory parameters also for M1 -> CB and PMC -> CB, as these were evident under both tDCS protocols. We focused the analysis on the Mid time window, based on the hypothesis that changes in connectivity parameters would be evident during behavioral and regional changes. Specifically, we submitted the modulatory parameters of Pu -> CB connection to a 2 × 2 rmANOVA with factors STIM (rCB tDCS, sham) and COND (SEQ, RND). We found a STIM x COND interaction (F_1,22_ = 4.5, p = 0.046) reflecting a reduction in negative modulation of Pu -> CB connection by SEQ during rCB tDCS compared to sham (t_22_ = 2.6, p = 0.02). No differences were observed for RND (p > 0.8, see Fig. 5C). There was no correlation between behavioral parameters of learning and modulatory effects during rCB tDCS (all p > 0.1).

### 3.7. M1 tDCS leads to increased negative modulation in putamen-cerebellar connection

Visual inspection of Fig. 5C suggested that lM1 tDCS affected modulation of Pu -> CB connection differently compared to sham. We therefore performed an exploratory analysis in Early and Late, using a 2 × 2 rmANOVA with factors STIM (lM1, sham) and COND (SEQ, RND). For Early, we found a significant COND x STIM interaction (F_1,22_ = 5.9, p = 0.02). Post-hoc t-tests suggest that this effect stems from stronger negative modulation during RND during lM1 tDCS compared to sham (t_22_ = 2.6, p = 0.02, see Fig. 5C). For Late, we found a main effect of STIM (F_1,22_ = 5.5, p = 0.03) but no interaction (p > 0.2) suggesting stronger negative modulation during lM1 tDCS compared to sham.

## Discussion

Whilst numerous behavioral studies have shown that tDCS can influence motor learning performance, only few have investigated the underlying network interactions that may drive these changes. In this study we addressed the neurophysiological effects of tDCS over M1 and cerebellum on learning-related activity, i.e. differences in activation between conditions, in motor networks. Behaviorally, we observed faster reaction times during cerebellar tDCS compared to sham in Mid-Late time windows. This effect was absent during M1 tDCS. Performance in a completion task was also improved following cerebellar tDCS compared to sham, for correct assured response, reflecting superior training effects persisting into the post-training period. Following M1 tDCS performance remained similar to sham, indicating absence of even an immediate after-effect. Recording fMRI simultaneously to tDCS and task performance allowed us to delineate the neurophysiological changes that accompany this behavioral effect. Indeed, we observed a learning-related increased activity in right M1, left cerebellum lobule VI, right IFG, and right IPL during cerebellar tDCS (compared to sham), and specifically at the time cerebellar tDCS led to improved learning. While no learning-related changes in activity were observed for M1 tDCS compared to sham on the group level, we did find an association between learning-related activity increase in right M1, and left middle frontal cortex and better sequence performance under M1 tDCS but not during sham. This suggests that the effect of M1 tDCS on activity in these regions is dependent on individual sequence performance. We then proceeded with analyzing the cortico-striato-cerebellar network, asking how improved learning during cerebellar tDCS relates to connectivity patterns. To this end, we employed DCM and constructed 60 different models comprising a network of M1, premotor cortex, SMA, putamen, and cerebellum. We found that cerebellar tDCS led to a significant decrease in learning-specific negative modulation of putamen to cerebellum connection, reflecting decreased inhibition of cerebellum to allow better learning. In summary, these results suggest that cerebellar tDCS has a physiological effect on interactions within the motor learning network which leads to improvements in motor learning.

### The effects of cerebellar and M1 anodal tDCS on learning

Participants learned the sequence under all stimulation protocols and sequence performance improved with time. Importantly, we found that cerebellar tDCS improved sequence performance at Mid and Late time windows compared to sham. These results were corroborated by an examination of sequence knowledge following the main experiment: cerebellar tDCS led to better knowledge of the sequence when compared to sham. The findings here are in accordance with previous work showing the beneficial effect of anodal cerebellar tDCS on motor sequence learning (Ferrucci et al., 2013; Shimizu et al., 2017) but in contrast to a recent report showing the opposite effect (Ballard et al., 2019). Note that in the latter study, a multi-channel montage was used with a net current of 2mA compared to our protocol with 1mA. Note as well that the authors excluded six subjects (out of 35) of their cohort who had low performance levels. Surprisingly and in contrast to previous evidence (Kantak et al., 2012; Krause et al., 2016; Nitsche et al., 2003; Saucedo Marquez et al., 2013; Stagg et al., 2011; Waters-Metenier et al., 2014), M1 tDCS had no effect during any time window on sequence learning, as indexed by reaction-time differences between sequence and random blocks. In addition, no differences in offline sequence knowledge was observed when compared to sham. A possible explanation to the absence of any learning improvements following M1 tDCS could be the stimulation electrode montage, selected based on computational modelling (supplementary materials, section 1). Most of the studies mentioned above used the common montage (supp. Fig. 1A) which led to far less focal stimulation of M1, thus potentially involving premotor areas, frontal and prefrontal regions. These regions could also have contributed to enhanced learning. Alternatively, the proximity of the electrodes in our M1 montage could have led to shunting of the current by the skin and soft tissue (Vöröslakos et al., 2018) thus not reaching M1. This explanation is however less likely as we observed both general changes to the BOLD signal following M1 tDCS, as well as association between the BOLD signal, in a number of motor regions, and sequence performance during M1 tDCS. In addition, recent evidence (Lefebvre et al., 2019) points to strong inter-individual variability of motor cortex excitability, when tDCS is placed on M1 (anatomical or “hotspot”) which could lead to lack of significant effects.

### The effect of tDCS on BOLD activity during motor learning

Specific learning-related activity changes were found for cerebellar tDCS only (compared to sham) and in the Mid time window, matching behavioral differences between cerebellar tDCS and sham. Specifically, cerebellar tDCS increased activity in left cerebellum lobule VI, right M1, right IPL and right IFG, during sequence relative to random blocks. These effects were absent in sham and in lM1 tDCS. These regions have been consistently shown to be modulated during motor learning (Hardwick, 2013) and are part of the cortico-striato-cerebellar network (Doyon 2009). Note that here too, the actually stimulated regions, left M1 and right anterior cerebellum did not show any learning-related changes compared to sham stimulation. Similarly, a concurrent continues theta burst stimulation and fMRI study investigating motor sequence learning (Steel et al., 2016), could not show any regional activation changes due to deactivation of M1. Instead, they observed reduced functional connectivity in visual and dorsal premotor cortex and increased connectivity within cingulate, and superior frontal gyrus. On the other hand, Waters and colleagues (2017) showed that anodal tDCS over M1 during training of a motor sequence led to specific increase over sensorimotor cortex during sequence performance when compared to sham. While our results showed no effect of M1 tDCS (compared to sham) on learning-related brain activity on the group level, we did find an association between sequence performance and increased learning-related activity in right M1 and left middle frontal cortex under M1 tDCS during the Late time window, which was significantly different than correlation coefficients in these regions under sham. This means that increased activity in these regions during sequence performance (compared to random) was dependent on how well subjects have learned the sequence at the end of the task. No such dependencies were found in other time windows or under cerebellar tDCS/sham. This result calls for a more individualized approach to target M1 using tDCS.

Why would tDCS lead to specific learning-related increase in activity in brain regions of the motor learning network? To answer this question, we have to gain better understanding of the effect of tDCS on neurophysiological parameters. Anodal DC applied in a M1 slice preparation and combined with low frequent synaptic activation leads to a long-lasting synaptic potentiation that requires NMDA receptor activation (Fritsch et al., 2010) and increased spine density (Gellner et al., 2020). Importantly, the authors showed that the effect of anodal direct current in M1 slices depends on activity-dependent BDNF secretion and this, in turn, has been shown to affect motor learning (Fritsch et al., 2010). Such physiological evidence is lacking for cerebellar tDCS, for which mechanisms may completely differ compared to neocortex stimulation. Despite that, it has been proposed that anodal tDCS to cerebellum increases the excitability of Purkinje cells which leads to stronger inhibitory output by deep cerebellar nuclei (DCN) and in turn to reduction in thalamic facilitation of cortical structures (Grimaldi et al., 2016). As sequence learning modulates activity in those regions and affects network interactions (Fletcher et al., 2005; Sun et al., 2007; Tzvi et al., 2015, 2014), it is plausible that together with cerebellar tDCS, differential increase in activity could be observed.

Only very few studies to date have conducted experiments combining simultaneous cerebellar tDCS and fMRI (D’Mello et al., 2017; Küper et al., 2019). One such study (Küper et al., 2019) explored the effect of cerebellar tDCS on the BOLD signal during simple finger tapping and did not observe any changes in activity following anodal stimulation, similar to our findings here on general motor-related activity (see supplemental materials, section 5). Note however, that the authors have focused their analysis on the cerebellum and did not assess possible differences due to cerebellar tDCS at the neocortex level. A different study explored the effect of anodal cerebellar tDCS on semantic processing (D’Mello et al., 2017). The authors likewise observed a differential increase in activity in language-related regions in a specific experimental condition (semantic prediction), similarly to our results. Against this background, we suggest that learning-related increased activity in right M1, left cerebellum, left IFG and right IPL, associated with better performance, reflect specific changes to the motor learning network following cerebellar tDCS.

### The effect of tDCS on interactions in the motor learning network

We hypothesized that tDCS would affect interactions within the motor learning network. Our imaging results showing activity increase during learning in M1, IFG, IPL and cerebellum following cerebellar tDCS, support the hypothesis that effects may be evident on a network level. We used DCM to explore the effect of tDCS on effective connectivity between a subset of regions comprising the cortico-striatal-cerebellar network. Comparing 60 different models, we found a different winning model for each stimulation protocol. Modulation of connection from M1 to cerebellum and PMC to cerebellum was evident in both tDCS protocols, whereas modulation of putamen to cerebellar connection was evident across all simulation protocols (including sham). This means that independent from tDCS, putamen to cerebellar connection is negatively modulated by the task, replicating out previous results (Tzvi et al., 2015). On the parametric level, we compared the modulatory effects of putamen to cerebellar connection across the different stimulation protocols. We found that cerebellar tDCS attenuated the modulation of this connection by SEQ in the Mid time window. This means that putamen strongly inhibits the cerebellum during sequence learning at sham, while cerebellar tDCS dampens this effect. Notably, this is a qualitative interpretation of these results, as the summed effect of the model on activity in the cerebellum is influenced by “default” intrinsic connections, the effect of the input on activity in the cerebellum (see supplementary materials, section 6 as well as supp. Tables 3 and 4) as well as other interactions in this network (i.e. negative modulation of M1 to cerebellum).

Neuroanatomical studies have shown a disynaptic pathway connecting the sensorimotor territory of putamen in macaque monkeys and the dentate nuclei in cerebellum, possibly through the thalamus (Bostan et al., 2010). This disynaptic pathway can assist fast communication between the cerebellum and the basal ganglia which may enable coordinated output to the motor cortex (Chen et al., 2014). Indeed, an experiment in mice showed that high-frequency stimulation of the cerebellar dentate nucleus led to an increased response in dorsolateral striatum. Bostan and Strick (2018) hypothesized that the integration of supervised, error-based, processes in the cerebellum and reinforcement mechanisms in putamen may underlie communication between these two structures during motor learning. The cerebellum receives an efference copy of an outgoing motor command and forms an internal model that predicts the movement outcome based on an updated error signal (Blakemore et al., 1998; Imamizu et al., 2000; Wolpert et al., 1998). As learning progresses and accuracy improves, these error signals decrease, and cerebellar activity as well. Application of tDCS to cerebellum led to increased cerebellar activity during learning which might have led to a stronger error-signal resulting in faster internal model formation compared to sham. Our results suggest that this process is influenced through an interaction with putamen: tDCS to cerebellum led to disinhibition of cerebellum by putamen during learning, possibly for communicating corrected motor commands to the basal ganglia, and then the motor cortex, leading to better learning. Note however that this effect might also be independent from learning as no correlations between modulatory parameters and behaviour were evident.

Interestingly, while this interpretation of the connectivity patterns is in accordance with increased activity in left cerebellum, it does not explain the increased learning-related activity in right M1, left IFG and right IPL. Our imaging results challenge the notion of cerebellar tDCS effects on cortical activity through inhibitory output of DCN by increased excitation of Purkinje cells on the cerebellar surface. This model however does not account for the effect of parallel and climbing fibers on synapses with Purkinje cells in DCN, also considered essential for controlling cerebellar-dependent learning (Ito, 2006). These may modulate the output of DCN, leading to increased putamen activity (see above) and as a result to increased cortical activity through the thalamus during learning.

### Limitations

Some limitations need to be considered. First, in terms of behavioral effects, we hypothesized improved learning performance due to both lM1 tDCS and rCB tDCS. We computed several post-hoc t-tests that were not corrected for multiple comparisons. This means that the reported results could be viewed as marginal in terms of their effect. This is true for the fMRI analyses as well in which tests were performed in three different time windows, without correction. In the DCM analysis, we focused on one connection (Pu -> CB) and one time window (Mid) for our analysis of rCB tDCS effects but the reported effects of lM1 tDCS were not corrected across the different time windows.

We found that subjects improved their learning of a new sequence with practice of a different sequence (across sessions). This presents a potential problem as this effect might interfere with observed changes due to the stimulation protocol. We therefore additionally tested using a between-subject design the effects of tDCS on learning (see supplementary materials, section 4) in each session separately using a mixed effects ANOVA. We found a COND x STIM x TIME interaction in the last session, which resulted from significant differences between cerebellar tDCS and sham, with no differences found for M1 tDCS. These results corroborate the original analysis showing an effect for cerebellar tDCS.

## Conclusions

We investigated the neurophysiological correlates of motor learning under transcranial direct current stimulation (tDCS). Our results show that right cerebellar tDCS led to improved sequence performance in Mid-Late time windows as indexed by decreased reaction times in the SRTT and improved assured sequence production in a completion task. Our concurrent imaging data shows learning-specific increase in activity during cerebellar tDCS in regions associated with the motor learning network. Network analysis with DCM revealed additional changes to putamen-cerebellar interactions under cerebellar tDCS that could lead to the learning benefits observed in our study. Together, these results show on the neurophysiological level, the specific effect of cerebellar tDCS on activity and connectivity in the motor learning network. Future work building on these results could establish network-driven models for post-stroke rehabilitation therapies using cerebellar stimulation.

## Acknowledgements

We would like to thank Gregor Spitta, Mourad Zoubir, Jennifer Nabel and Susanne Schellbach for their assistance data collection and David Minker for his assistance with experimental design. We are also grateful for three anonymous reviewers for their valuable comments and suggestions. This work is supported by the DFG grant TZ 85/1-1 to ET.

## List of abbreviations

tDCS: transcranial direct current stimulation
M1: primary motor cortex
CB: cerebellum
PMC: premotor cortex
SMA: supplementary motor area
Pu: putamen
DCN: deep cerebellar nuclei
IPL: inferior parietal lobe
IFG: inferior frontal gyrus
fMRI: functional magnetic resonance imaging
RT: reaction times
DCM: dynamic causal modelling
SEQ: sequence
RND: random
SMP: simple
SRTT: serial reaction time task
BOLD: blood oxygen level dependent
MNI: Montreal Neurological Institute
GLM: general linear model
COND: condition
STIM: stimulation
BMA: Bayesian model averaging
rmANOVA: repeated measures analysis of variance;

## Notes

### Competing Interest Statement

The authors have declared no competing interest.

