## supplementary materials for "Beneficial effects of cerebellar tDCS on motor learning are associated with altered putamen-cerebellar connectivity: a simultaneous tDCS-fMRI study"

1. Computational modelling of M1 and cerebellar-stimulation locations

We used the SimNIBS 2.1 software package (Saturnino et al., 2019) to simulate two different electrode montages in order to test for optimal electrode placement (Thielscher et al., 2015). The head model, made available by the software package, was created using finite element modeling on T1- and T2-weighted MRI images of an exemplary subject resulting in a high-resolution tetrahedral head mesh model containing 8 tissue types. We set the conductivities of brain white matter, gray matter and of cerebrospinal fluid to 0.126 S/m, 0.275 S/m and 1.654 S/m respectively (Opitz et al., 2015). Values of 0.025 S/m and 0.008 S/m were used for the spongy and compact bone of the skull. All tissues were treated as isotropic. The electrical field $\vec{E}$ was determined by taking the numerical gradient of the electric potential. For both montages we used 3 cm x 3 cm rectangular electrodes with 3 mm electrode thickness and 2 mm paste thickness. The total current injected was 1 mA.

In the standard M1 montage, used primarily in previous work to stimulate M1, the anode was placed over left M1 and the cathode over right supraorbital ridge (supp. Fig. 1A). We tested an alternative montage by placing the same electrodes directly anterior and posterior to M1 (Fig. 1B). The simulations show that the alternative montage presented in Fig. 1B, results in a strong focal distribution of the electric field over M1, whereas the standard montage (supp. Fig. 1A) results in a strong electric field over M1, premotor as well as contralateral prefrontal cortex.

For simulating cerebellar tDCS, we used as starting point a montage (supp. Fig. 1C) in which the anodal electrode was placed 3 cm right lateral to the inion and the reference (cathodal) electrode over the right buccinator muscle (van Dun et al., 2016). Then, we tested a reference for cerebellar stimulation montages on the neck (Supp. Fig. 1E), as well as a reference over right mandibula (Fig. 1B). In all cerebellar montages the anode was kept 3 cm right lateral to the inion. All parameters were kept identical to the M1 montage, including the size of stimulation electrodes. The montage with the reference over right buccinator muscle (supp.

Fig. 1C) showed enhanced electric field distribution over temporal cortex, which could affect memory processes relevant to our task. The montage with the reference placed over the neck (supp. Fig. 1E) showed enhanced electric field distribution in the brainstem which may lead to physiological adverse effects. According to this simulation and the given models, placing the reference over right mandibula thus seems to be the optimal montage with more focal effects in the cerebellum and less current distribution over temporal cortex and brainstem. Given these results, we decided to place electrodes directly anterior and posterior to left M1 for left M1 tDCS and lateral to the inion as well as over right mandibula for right cerebellar tDCS (Fig. 1B).

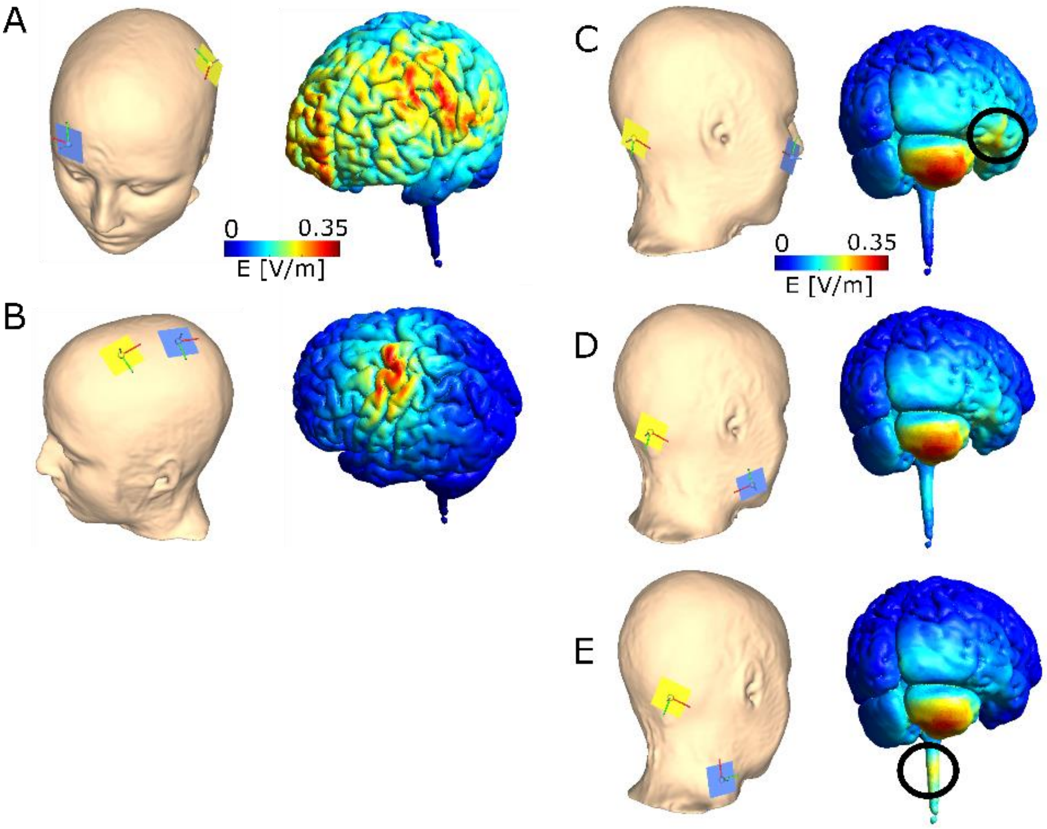


***Supplementary Figure 1.*** Electric field distribution was estimated for different tDCS montages. **A** M1 standard montage with anode over left M1 and cathode over right supraorbital ridge **B** M1 montage with anode anterior- and cathode posterior to the central sulcus. Cerebellar standard montage with anode over cerebellum (3 cm right lateral to the inion, **C-E**) and cathode over ipsilateral buccinators muscle (**C**), over the mandibula (**D**), or on the neck (**E**). The black circles show the areas with increased electric field outside of the cerebellum.

### 2. <u>Behavioral results: motivation, tiredness, sham blindness, adverse effects</u>

At the beginning of each experimental session, subjects reported their level of fatigue and their motivation for participation (both on a 1-10 Likert scale). At the end of each experimental session, subjects reported their fatigue level again as well as whether the stimulation was painful, and whether they thought it was real or sham stimulation.

Motivation to engage in the experiment was very high overall (mean = 8.5/10) and did not differ between sham, rCB and lM1 tDCS (all $p > 0.6$). There was no difference in fatigue levels prior to stimulation between the experimental sessions (all $p > 0.5$). Moreover, the difference in tiredness before and after stimulation did not differ between sham, rCB and lM1 tDCS (all $p > 0.8$). After subjects completed the experiment, we asked them to state whether they had undergone sham or real stimulation. In 5 out of 25 cases of sham, subjects stated correctly that they received sham stimulation (20%). The chi-square test determined that this proportion is even significantly lower than chance level (50%, $p = 0.03$, $\chi^2 = 4.95$). For real tDCS, subjects stated wrongly in 10 out of 50 cases that they did not receive brain stimulation (20%). Here as well the chi-square test showed that this proportions is far below chance (50%, $p = 0.002$, $\chi^2 = 9.89$). This shows that our stimulation protocol worked and that participants could not distinguish between sham and real stimulation.

We also asked the subjects whether they experienced phosphenes which would be a sign of visual cortex stimulation (Kar and Krekelberg, 2012). One subject reported phosphenes during cerebellar tDCS and one during sham over cerebellum. A total of six subjects reported light pain in the back of the head or neck during tDCS. In five subjects the report followed cerebellar tDCS and in one following sham. One subject reported dizziness following cerebellar tDCS. One other subject reported somatosensory sensation following cerebellar tDCS. None of the reports influenced performance in the task (according to self-evaluation by the subjects).

3. <u>Behavioral results: Analysis of RTs in RND blocks of the SRTT</u>

We analyzed RTs of RND blocks as it seems participants were slower in the second mini-block (Mini2) compared to the first mini-block (Mini1) and that this effect increases as learning progresses. Here as well, we additionally explored whether stimulation of lM1 or rCB has an effect on slowing-down during performance of RND trials. We therefore subjected the RTs of RND mini-blocks only to a rmANOVA with factors STIM (rCB, lM1, sham), MiniBlock (Mini1, Mini2) and TIME (time points 1-5). A main effect of MiniBlock ($F_{1,23} = 64.5$, $p < 0.0001$), a main effect of TIME ($F_{4,92} = 5.7$, $p < 0.001$) as well as MiniBlock x TIME interaction ($F_{3,72} = 8.2$, $p < 0.001$) demonstrating that the slowing down in Mini2 compared to Mini1 is indeed increasing with time and could be relevant for learning processes. LM1 or rCB tDCS has however no effect on learning processes reflected in RT slowing in RND mini-blocks, evident by lack of STIM x MiniBlock or STIM x MiniBlock x TIME interactions ($p > 0.7$).

4. Behavioral results: Analysis of learning improvement across sessions

As subjects performed the SRTT a total of three times across three sessions, we also tested whether sequence learning significantly improved across sessions using rmANOVA, with factors SES (session order 1-3), COND (SEQ, RND) and TIME (Early, Mid, Late) which revealed a main effect of SES ($F_{2,46} = 9.3$, $p < 0.001$), a COND x SES ($F_{2,46} = 4.4$, $p = 0.02$) interaction and a COND x SES x TIME ($F_{4,92} = 4.7$, $p = 0.002$) interaction suggesting that indeed subjects could learn better the more times they performed the task although each time a different sequence was used. To find whether this session effect was driving the differences we found between rCB tDCS and sham, we compared in each session between the different groups that received either rCB tDCS, lM1 tDCS and sham using a mixed effects ANOVA accounting for between-subjects STIM factor and within-subjects COND (SEQ, RND) and TIME (Early, Mid, Late) factors. In the last session (lM1 = 8 subjects, rCB = 6 subjects, sham = 10 subjects) we found COND x STIM x TIME interaction ($F_{4,42} = 2.9$, $p = 0.04$) suggesting that the stimulation effects were not driven by learning carry-over effects. Post-hoc tests (Wilcoxon rank sum, correcting for multiple comparisons) on SEQ-RND differences at each transfer from RND to SEQ showed significant differences between rCB tDCS and sham in the last session at the second transfer in Mid time window ($p = 0.005$), and at the second transfer in Late time window ($p < 0.001$). No differences were found between lM1 tDCS and sham, as well as between lM1 and rCB tDCS. This analysis suggests that improved learning due to rCB tDCS is mostly driven from session 3 in this task, in which subjects could reach a learning plateau.

5. Imaging results: the effect of tDCS location on motor-related activity

We investigated task-related changes in brain activity comparing lM1 to rCB tDCS across the different time windows (Early, Mid and Late), using a flexible factorial design with factors STIM (lM1, rCB) and COND (SEQ, RND) in each of these time windows, analog to the analyses in the main text comparing tDCS to sham. In Mid, we found enhanced activity during lM1 compared to rCB tDCS in right CB crus I and crus II (supp. Fig. 3), bilateral medial superior frontal gyrus and left SMA (supp. Table 2). In Late, enhanced activity during lM1 compared to rCB tDCS was evident in left precuneus and cuneus (supp. Table 2). Together, these results suggest that lM1 tDCS during motor task performance leads to a wide-spread activity increase in motor networks.

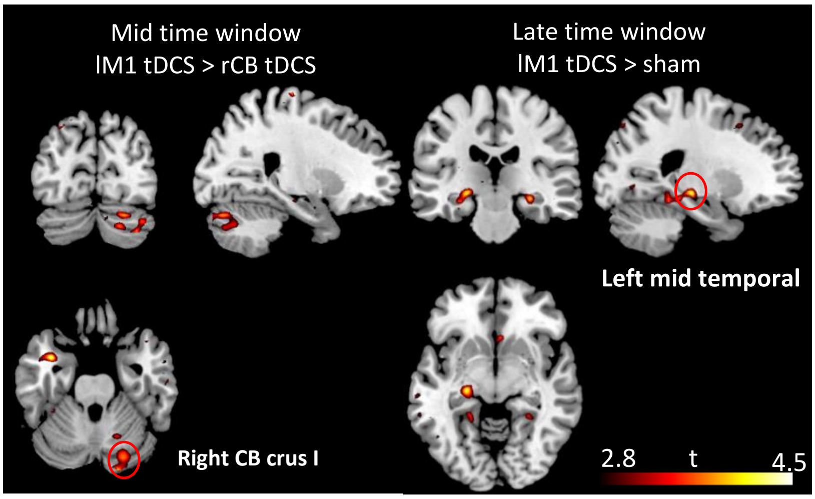


***Supplementary Figure 3.*** General effects of tDCS on activity during motor performance. LM1 tDCS resulted in increased activations in a Mid time window compared to rCB tDCS, and in a late time window compared to sham.
All activations are shown on a $p < 0.001$ voxel-level threshold with $p < 0.05$, family-wise error corrected on the cluster level for multiple comparisons.

6. <u>DCM results: endogenous connections in each of the optimal models</u>

The following analyses were focused on endogenous effects on connections in the optimal models. We tested for significant endogenous connections for each stimulation protocol. The results are presented in supp. Table 3. We considered endogenous connections as significant with a Bonferroni corrected p-value across the 20 connections (0.05/20 = 0.0025). Notably, we found positive endogenous connections from CB to M1, SMA and PMC across all stimulation modalities. All endogenous connections from CB to other structures were positive as well (except for M1 stim in connection from CB to Pu). In addition, connections from M1 to CB and from Pu to SMA, were positive for rCB tDCS only. Some negative connections were evident by trend ($p < 0.05$, uncorrected, see supp. Table 3).

We then explored whether endogenous connections were affected by rCB or by lM1 tDCS compared to sham. Note that since each stimulation protocol had a different winning model, we used Bayesian model averaging to compute a weighted average of parameter estimates across all models. No differences between lM1 tDCS or rCB tDCS and sham were observed (all $p > 0.1$).

7. <u>DCM analysis of bilateral M1-cerebellar network</u>

We defined a 4-region network including left and right M1 as well as left and right cerebellum. Time series from left M1 and right cerebellar VOIs are described in the main text. Similarly, time series for right M1 (48,-30,60) and left cerebellum (-38,-58,-34) VOIs were extracted. We specified 27 models, all with input to right cerebellum (rCB, similarly to the analysis in the main text) and fully connected VOIs. Modulatory effects were hypothesized to occur either between bilateral M1, bilateral CB or in rM1-lCB as well as lM1-rCB connections (see supp. Fig. 4 below).

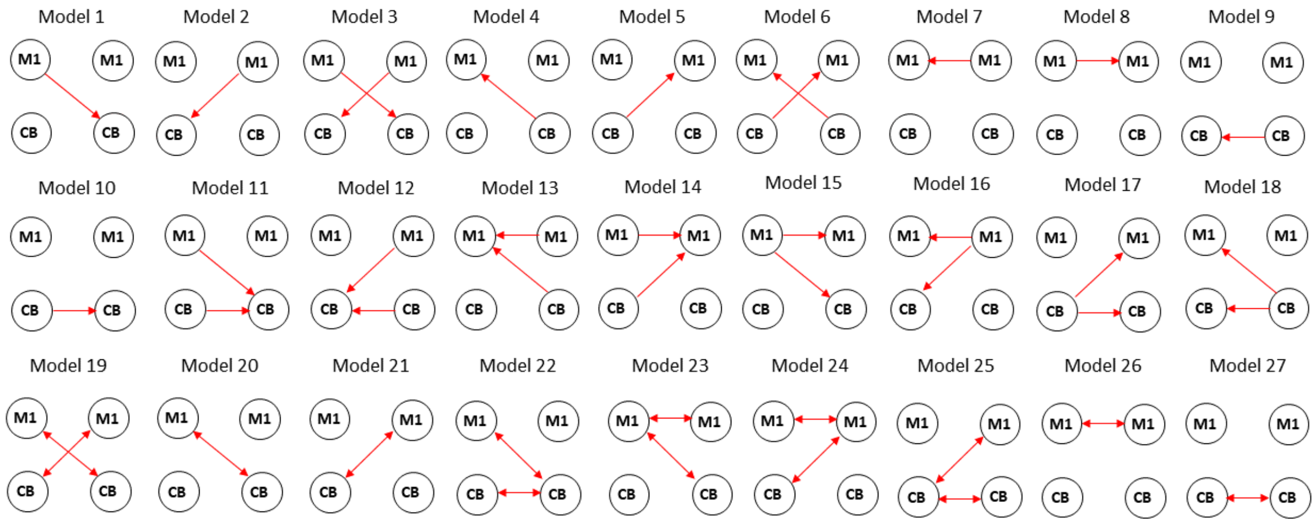


**Supplemental Figure 4:** model space for dynamic causal modeling analysis. The arrows mark the connections that were assumed to be modulated by the task. Note that all VOIs were fully

Bayesian model selection procedure with a random-effects model was performed for each of the stimulation protocols separately, in order to account for changes in network architecture due to tDCS. Under all stimulation protocols, the optimal model was model 24, which had modulation of bilateral M1, and M1-CB connections (Exceedance probability: rCB tDCS: p_ex = 0.224; lM1 tDCS: p_ex = 0.23; sham: p_ex = 0.56). For rCB tDCS, the next probable model, model 18 had modulation of rCB to both lCB and left M1 connection with a very close exceedance probability (p_ex = 0.217). Indeed, the probability of equal model frequencies was p = 0.72.

Next, we used the script spm_dcm_fMRI_check.m to ensure that inversion of the winning model converged. The script estimates the percentage variance explained of the winning model in all subjects and sessions. We found that the winning model did not converge in many subjects (variance explained < 10%: lM1 tDCS 9/23; rCB tDCS 15/23; sham: 16/23). We therefore were not able to draw statistically meaningful conclusions from the parameters of the winning model across stimulation protocols.

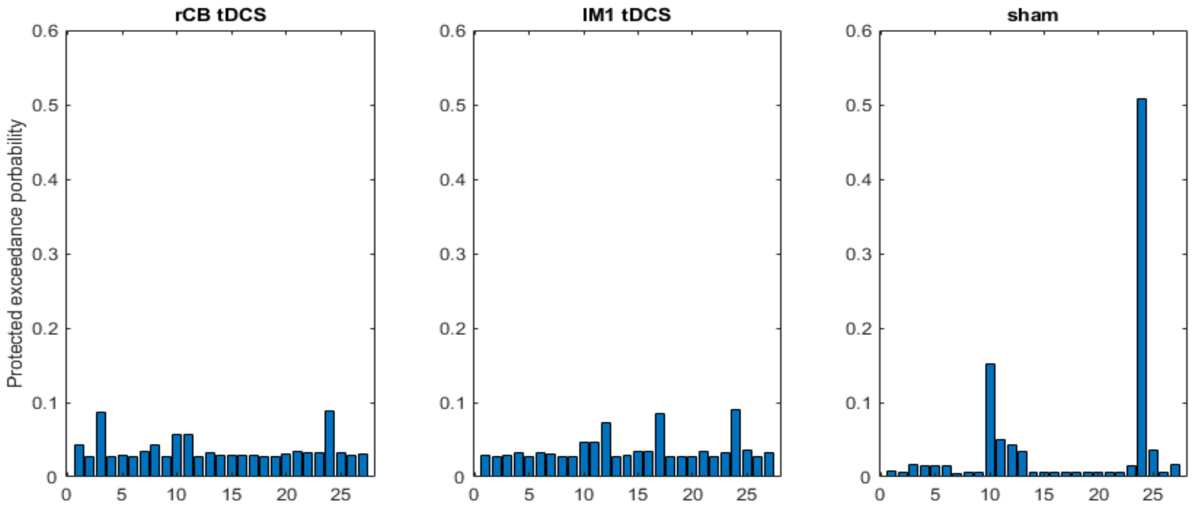


**Supplemental Figure 5:** Results of Bayesian model selection across the 27 models (described in supplemental Figure 4) for each of the stimulation protocols.

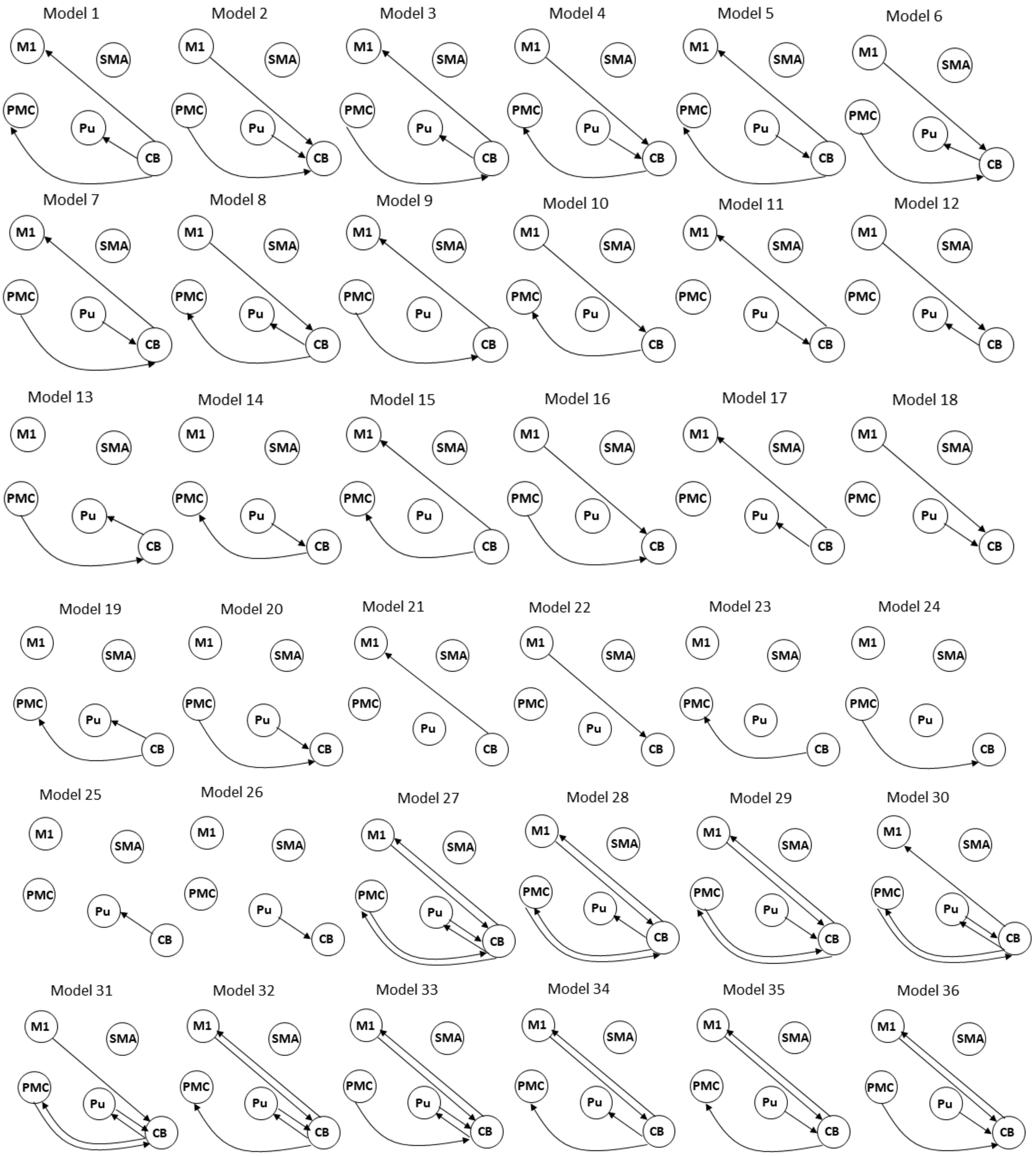
Model 1
Model 2
Model 3
Model 4
Model 5
Model 6
Model 7
Model 8
Model 9
Model 10
Model 11
Model 12
Model 13
Model 14
Model 15
Model 16
Model 17
Model 18
Model 19
Model 20
Model 21
Model 22
Model 23
Model 24
Model 25
Model 26
Model 27
Model 28
Model 29
Model 30
Model 31
Model 32
Model 33
Model 34
Model 35
Model 36
M1
SMA
PMC
Pu
CB

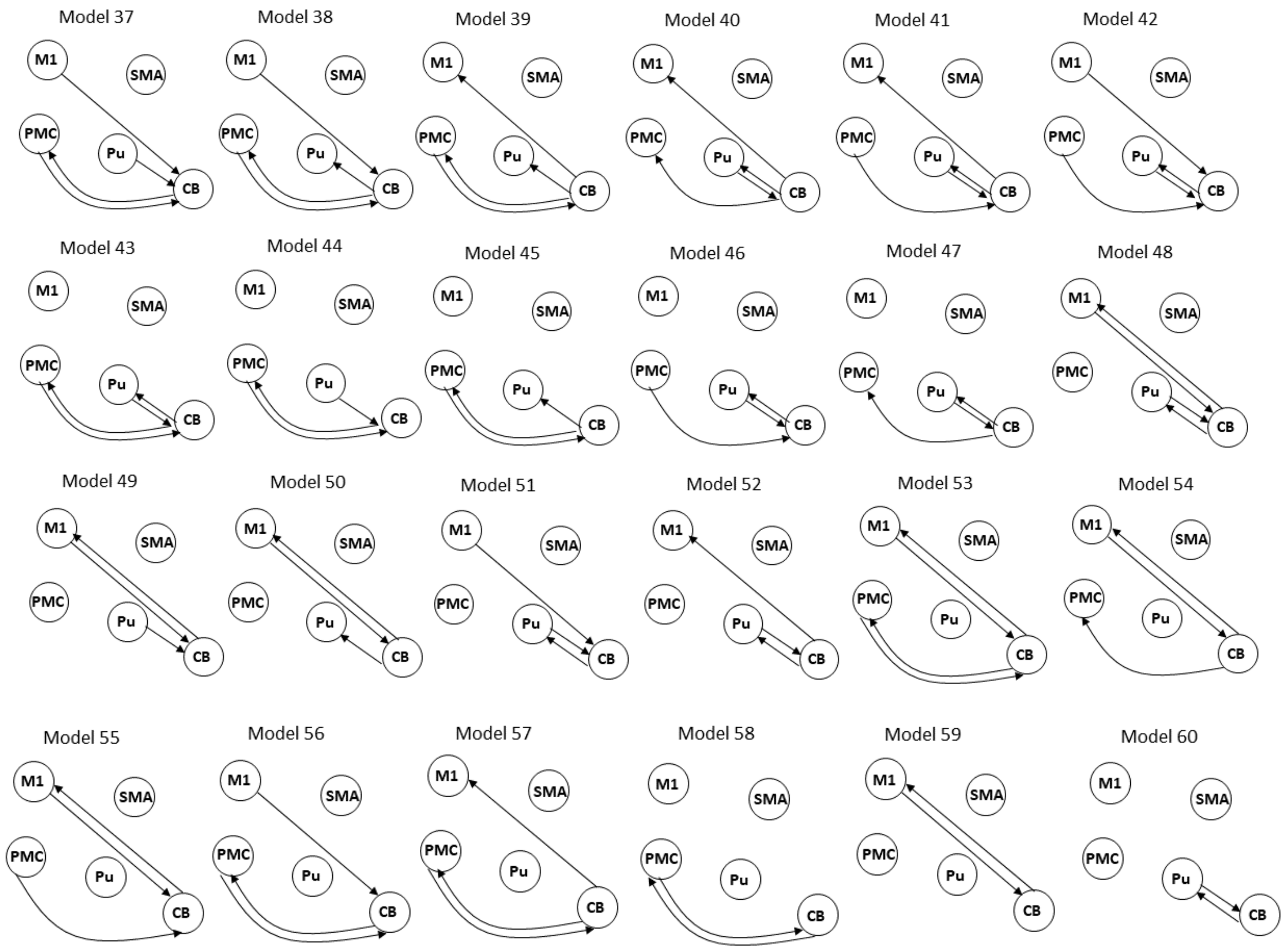


**Supplementary Figure 2.** Model space for dynamic causal modelling analysis. The arrows mark the connections in each model which were assumed to be modulated by the task. Note that all VOIs were fully connected (not shown).

**Supplementary Table 1: regions of interest for DCM analysis**

| Region | MNI-coordinates | t-value (lM1 tDCS, rCB tDCS, sham) |
|---|---|---|
| lM1 | -40 -26 56 | 10.1, 12.3, 13.0 |
| lSMA | -4 -2 56 | 9.0, 10.0, 14.3 |
| lPMC | -30 -16 62 | 9.5, 8.4, 9.6 |
| lPu | -24 -4 8 | 7.7, 7.1, 7.2 |
| rCB | 20 -50 -22 | 9.9, 9.1, 13.1 |

**Supplementary Table 2: fMRI task activations ($p < 0.05$, cluster-level FWE)**

| Region | MNI-coordinates | | | t-value | n-voxels |
|---|---|---|---|---|---|
| lM1 tDCS > rCB tDCS: Main effect STIM (Mid) | | | | | |
| R cerebellum crus I | 18 | -88 | -26 | 4.20 | 108 |
| R cerebellum crus II | 32 | -80 | -40 | 4.09 | 59 |
| R medial sup. frontal gyrus | 2 | 26 | 60 | 4.09 | 50 |
| L medial sup. frontal gyrus | -6 | 24 | 62 | 3.88 | 51 |
| L supplementary motor area | -4 | 22 | 60 | 3.23 | 21 |
| lM1 tDCS > rCB tDCS: Main effect STIM (Late) | | | | | |
| L precuneus | -12 | -58 | 28 | 4.10 | 107 |
| L cuneus | 2 | -78 | 24 | 3.97 | 134 |
| lM1 tDCS > sham: Main effect STIM (Late) | | | | | |
| L middle occipital gyrus | -32 | -72 | 32 | 4.27 | 153 |
| L Angular gyrus | -42 | -68 | 30 | 3.92 | 64 |
| L middle temporal gyrus | -46 | -64 | 20 | 3.54 | 25 |

**Supplementary Table 3: posterior estimates of endogenous connections in the winning model (mean±SE)**

| Connection | M1 tDCS | | CB tDCS | | Sham | |
|---|---|---|---|---|---|---|
| | Ep (Hz) | p-val | Ep (Hz) | p-val | Ep (Hz) | p-val |

| | | | | | | | |
|---|---|---|---|---|---|---|---|
| lM1 -> | rCB | 0.15±0.05 | 0.003 | **0.27±0.07** | **0.001** | 0.14±0.06 | 0.02 |
| lM1 -> | lPu | -0.05±0.04 | 0.21 | -0.04±0.04 | 0.29 | 0.01±0.05 | 0.79 |
| lM1 -> | lSMA | 0.05±0.04 | 0.23 | -0.03±0.05 | 0.65 | 0.12±0.06 | 0.05 |
| lM1 -> | lPMC | 0.03±0.04 | 0.39 | 0.03±0.03 | 0.37 | 0.08±0.04 | 0.03 |
| rCB -> | lM1 | **0.47±0.05** | **<0.001** | **0.46±0.05** | **<0.001** | **0.46±0.06** | **<0.001** |
| rCB -> | lPu | 0.10±0.05 | 0.04 | **0.18±0.04** | **<0.001** | **0.12±0.03** | **<0.001** |
| rCB -> | lSMA | **0.29±0.05** | **<0.001** | **0.34±0.04** | **<0.001** | **0.31±0.05** | **<0.001** |
| rCB -> | lPMC | **0.35±0.05** | **<0.001** | **0.33±0.04** | **<0.001** | **0.35±0.04** | **<0.001** |
| lPu -> | lM1 | -0.08±0.05 | 0.14 | -0.13±0.04 | 0.009 | -0.12±0.06 | 0.04 |
| lPu -> | rCB | -0.10±0.06 | 0.15 | -0.12±0.08 | 0.15 | -0.19±0.07 | 0.009 |
| lPu -> | lSMA | 0.11±0.03 | 0.004 | **0.15±0.04** | **0.001** | 0.08±0.04 | 0.05 |
| lPu -> | lPMC | -0.03±0.04 | 0.39 | -0.02±0.03 | 0.44 | -0.11±0.04 | 0.009 |
| lSMA -> | lM1 | -0.07±0.05 | 0.16 | -0.09±0.06 | 0.15 | -0.09±0.05 | 0.10 |
| lSMA -> | rCB | -0.14±0.06 | 0.02 | -0.11±0.05 | 0.03 | -0.09±0.05 | 0.10 |
| lSMA -> | lPu | 0.09±0.05 | 0.12 | 0.10±0.03 | 0.007 | 0.13±0.06 | 0.03 |
| lSMA -> | lPMC | -0.06±0.05 | 0.34 | -0.00±0.04 | 0.9 | -0.06±0.02 | 0.03 |
| lPMC -> | lM1 | 0.15±0.05 | 0.01 | 0.07±0.04 | 0.06 | 0.07±0.03 | 0.04 |
| lPMC -> | rCB | 0.12±0.06 | 0.07 | **0.17±0.05** | **0.001** | **0.17±0.05** | **0.002** |
| lPMC -> | lPu | 0.07±0.04 | 0.11 | 0.05±0.03 | 0.16 | -0.03±0.04 | 0.44 |
| lPMC -> | lSMA | 0.10±0.05 | 0.04 | 0.05±0.03 | 0.16 | -0.01±0.04 | 0.84 |

*Bonferroni corrected for multiple comparisons: p = 0.0025

**<u>Supplementary Table 4: posterior estimates of the input to cerebellum in the winning model (mean±SE)</u>**

| | **M1 tDCS** | | **CB tDCS** | | **Sham** | |
|---|---|---|---|---|---|---|
| | SEQ | RND | SEQ | RND | SEQ | RND |
| Early | 0.19±0.02 | 0.28±0.02 | 0.15±0.01 | 0.24±0.02 | 0.25±0.03 | 0.23±0.03 |
| Mid | 0.25±0.02 | 0.23±0.02 | 0.19±0.02 | 0.21±0.02 | 0.22±0.02 | 0.21±0.02 |
| Late | 0.26±0.02 | 0.20±0.01 | 0.22±0.02 | 0.20±0.02 | 0.20±0.01 | 0.16±0.01 |
